# Role of protein-protein interactions on model chromatin organization

**DOI:** 10.1101/2024.03.03.583162

**Authors:** Pinaki Swain, Sandeep Choubey, Satyavani Vemparala

## Abstract

The three-dimensional organization of chromatin is influenced by DNA-binding proteins, through specific and non-specific interactions. However, the role of DNA sequence and interaction between binding proteins in influencing chromatin structure is not yet fully understood. By employing a simple polymer-based model of chromatin, that explicitly considers sequence-dependent binding of proteins to DNA and protein-protein interactions, we elucidate a mechanism for chromatin organization. We find that: (1) Tuning of protein-protein interaction and protein concentration is sufficient to either promote or inhibit the compartmentalization of chromatin. (2) The presence of chromatin acts as a nucleating site for the condensation of the proteins at a density lower than in isolated protein systems. (3) The exponents describing the spatial distance between the different parts of the chromatin, and their contact probabilities are strongly influenced by both sequence and the protein-protein attraction. Our findings have the potential application of re-interpreting data obtained from various chromosome conformation capture technologies, thereby laying the groundwork for advancing our understanding of chromatin organization.

## I. INTRODUCTION

Eukaryotic chromatin undergoes intricate spatiotemporal organization within the cell nucleus through three-dimensional compaction. Such spatial arrangement is crucial for regulating a variety of cellular processes, such as DNA replication, DNA damage repair, and gene expression. Chromatin is hierarchically organized across different length scales^1^, ranging from wrapping DNA around nucleosomes to the more complex higher-order folding of chromatin fibers into specific structural domains. Such structural domains include the transcriptionally active euchromatin (A) and the transcriptionally inactive heterochromatin (B). Additionally, advancements in various chromosome conformation capture technologies, such as Hi-C, coupled with live cell imaging tools, have unveiled a notably intricate picture of chromatin structure. At a scale of approximately 1*−* 10 Mbp, the AB compartments are discernible^2^, while at a smaller scale of 100 Kbp-1 Mbp, topologically associated domains (TADs) become apparent^3–5^. Furthermore, micro-C techniques, which capture chromosome folding at nucleosome-level resolution, have identified smaller loops of size 2 *−*10 Kbp^6^ and micro-compartments with AB-like architecture^7,8^. The underlying physical principles that dictate the formation of such nested organization of chromatin compartments and loops are still not fully understood.

Theoretically, various polymer-based models have been employed to elucidate aspects of chromatin’s spatial organization. These models broadly fall into two categories: loop-extrusion models and phase separation models of copolymers^9–12^. Loop-extrusion models effectively explain TAD formation^13–18^. In contrast, copolymer models have been employed to explain the formation of chromatin compartments and the promoter-enhancer interactions^8,19–24^. Block copolymers, comprising of two different monomer subchains, exhibit a layered organization in a poor solvent, a process known as microphase separation^25–29^ or micelle formation^30^. Microphase separation of block copolymers has also been used to explain phases of disordered proteins with hydrophobicity patterning^31^, localization of RNA segments inside paraspeckles^32^, and internal organization of biomolecular condensates^33–35^. However, it’s worth noting that these models often oversimplify or disregard the impact of protein binding to DNA and protein-protein interactions on chromatin organization. Interestingly, recent experimental studies have shown that protein-protein and protein-DNA interactions can lead to the formation ‘microphases’ on DNA as well as protein-DNA co-condensates. These mechanisms lead to condensate formation below the saturation concentration for bulk phase separation^36–40^. For instance, a recent *in vitro* study demonstrated that the pioneer transcription factor KLF4 can form surface condensates on DNA in a sequence-dependent manner through a prewetting-like transition^39^. Additionally, other studies have shown that pioneer transcription factor FoxA1 and heterochromatin protein 1 (HP1) can facilitate DNA compaction through protein-DNA and protein-protein interactions. Such co-condensation of protein and DNA is also known as polymer-assisted condensation (PAC)^40^. Therefore, it is crucial for polymer-based models of chromatin to consider the effect of sequence-dependent protein-DNA binding and protein-protein interactions for a comprehensive understanding of their role in chromatin structure and dynamics.

Over a decade ago, Nicodemi and colleagues introduced the ‘strings and binders switch’ (SBS) model, a unique approach to modeling chromatin that incorporates binding of proteins to DNA^41,42^. This model predicted chromatin condensation as a function of the concentration of binding proteins, a phenomenon also identified as bridging-induced phase separation (BIPS)^43–47^. Notably, SBS model has been frequently employed to calculate two experimentally reported quantities: the genomic loci exponent (*ν*), which describes how the average spatial distance between two chromatin segments scales with their contour distance ( ⟨*R*(*s*) ⟩ *∼s*^*ν*^ ), and the contact probability exponent (*α*), which quantifies the average frequency of contact between two distant segments ( ⟨*P*_*c*_(*s*) ⟩ *∼s*^*−α*^). The predicted values for the parameter *ν* range from 0 to 0.6, while *α* ranges from 0 to 2. The range for *ν* in the SBS model includes values predicted by the fractal-globule model (*ν* = 0.33 and *α* = 1.08) as a specific case^48,49^. However, most experiments report *ν* values between 0.3 and 0.5^50^, with *ν* = 0.1 as an exception, observed in FISH experiments by Jhunjhunwala et al., to which the SBS model compares its predictions^42,51^. Nonetheless, the SBS model does not consider interactions among the DNA binding proteins. A key question arises: can the incorporation of protein-protein interactions influence the experimentally measured range of genomic loci exponent (*ν*) and the contact probability exponent (*α*)?

Here we examine a simple polymer-based model of chromatin that explicitly accounts for DNA sequence-dependent binding of proteins and protein-protein interaction. To this end, we structure the paper as follows: In Section II A and Section II B, we detail the model and methods used in our study. Section III A focuses on the impact of incorporating explicit proteinprotein interactions and protein concentration into a homopolymeric chromatin model (-AAAAAA-). Our objective is to delineate the regimes of Bridging-Induced Phase Separation (BIPS), Polymer-Assisted Condensation (PAC), and Liquid-Liquid Phase Separation (LLPS). In Section III B, we explore how protein-protein interactions influence the genomic loci exponent (*ν*) and the contact probability exponent (*α*). Section III C examines the interplay between protein-protein interactions and sequence heterogeneity in determining chromatin compartmentalization. For this purpose, we expand our study to include two additional polymer sequences: a block copolymer (-AAA-BBB-) and an alternating polymer (- ABABAB-). This approach diverges from existing models, which typically assign different interactions among monomers to differentiate between heterochromatin and euchromatin^20,21,41,52,53^. In our model, we consider the chromatin polymer as a homogeneous self-avoiding polymer from the monomer perspective. However, heterogeneity arises from protein interactions: Monomers A and B have distinct binding affinities for proteins P. Throughout our simulations, the protein-chromatin interaction is consistently attractive and fixed, while we vary the protein-protein interactions and protein density. Our simulations reveal that the block copolymer sequence spontaneously forms a layered organization, unlike its alternative sequence counterpart.

## II. MODELING AND METHODS

### A. Model

Our model consists of a coarse-grained polymer representing chromatin and binders representing protein molecules (Fig. 1). Our model includes a homopolymeric chromatin composed entirely of monomer type A, and two heteropolymeric forms: a block copolymer and an alternating polymer, both consisting of two types of monomers, A and B. The model chromatin consists of *N* = 2000 monomers and the adjacent monomers in the polymer are bonded with harmonic potential

**FIG. 1.**
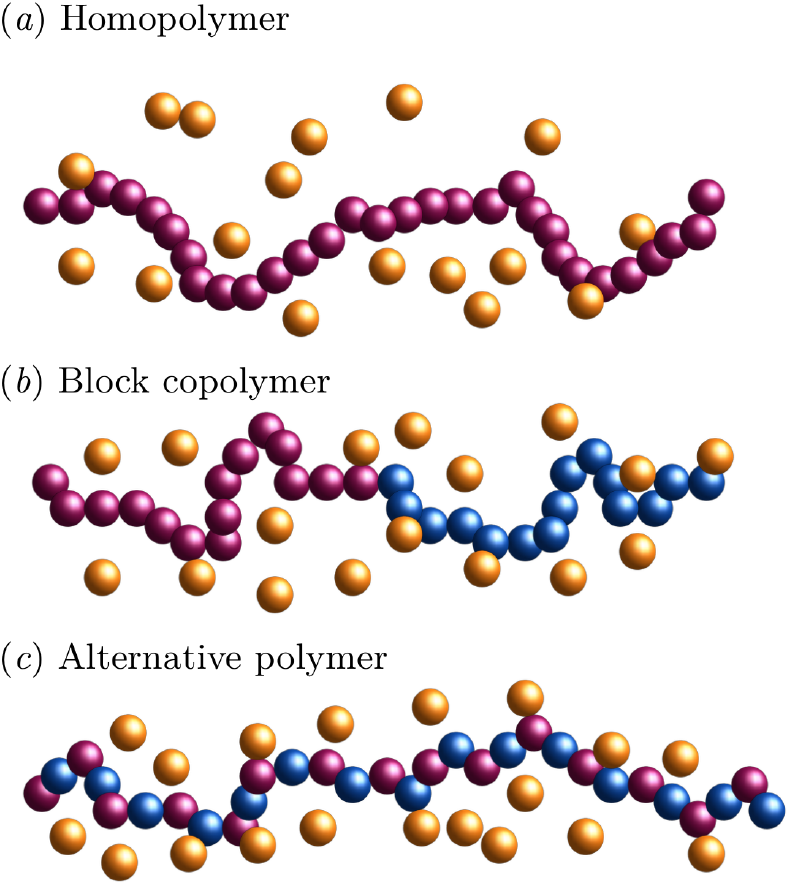
Schematic of different chromatin models: (*a*) homopolymer, (*b*) block copolymer, and (*c*) alternative polymer. The proteins, shown in yellow (P), bind to the magenta monomers (A) with an attractive strength of 1.5 *k*_*B*_*T* and bind to the cyan monomers (B) with a stronger attractive strength of 2 *k*_*B*_*T* . In all the cases, the proteins also interact between themselves with a strength of *ε*_*PP*_ , which we vary from 0.1 *k*_*B*_*T* to 2 *k*_*B*_*T* . We vary the protein density (*ρ*_*P*_ ) from 0.0005 to 0.006 *σ*^*−*3^ which translates to a protein concentration of 1.3 nM to 16 nM (see Sec. II A).

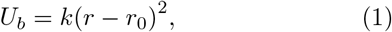

where, *k* = 100 *k*_*B*_*T/σ*^2^ is the bond energy and *r*_0_ = *σ* is the equilibrium bond length. Here, *k*_*B*_*T* is the thermal energy, and *σ* is the size of each monomer. In our model, the monomers and proteins are of equal size. In our computational model, non-bonded interactions between monomer-monomer, monomer-protein, and protein-protein pairs are simulated using the Lennard-Jones potential. However, for interactions amongst monomers within all three polymer configurations, we employ the Weeks-Chandler-Andersen (WCA) potential, which is purely repulsive^54^.

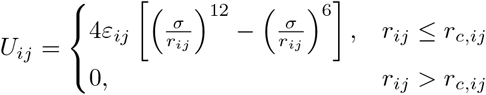

The interaction parameters, namely the strength (*ε*_*ij*_) and effective range (*r*_*c*,*ij*_) of the potential *U*_*ij*_, are specifically tailored for the diverse interacting pairs within our system. For instance, monomers of type A engage with proteins (notated as P) with an interaction strength of 1.5 *k*_*B*_*T* , denoting a moderate affinity. In contrast, type B monomers present a higher affinity towards these proteins, engaging with an interaction strength of 2 *k*_*B*_*T* . This differential in interaction strength is a simplified representation of the specificity and varied affinity observed within biological chromatin regions towards binding proteins. Additionally, the proteins themselves manifest mutual attraction, characterized by an interaction strength denoted as *ε*_*P* *P*_ . This feature enables the simulation of a spectrum of aggregation and phase separation phenomena, contingent upon protein concentration and interaction dynamics. It is important to underscore that within the heteropolymeric chromatin models (block and alternating polymers), the distinction between monomer types A and B is confined exclusively to their interactions with binding proteins. Beyond these specific interactions, the monomers are considered homogenous, sharing identical physical properties. This nuanced approach is pivotal for isolating the effects of protein interactions on chromatin behavior, circumventing the introduction of superfluous complexity from the chromatin architecture itself. Comprehensive details regarding all non-bonded interaction parameters operational within our system are systematically cataloged in Table II A.

**TABLE I.** Non-bonded interaction parameters for a model chromatin with two kinds of monomers, A and B, in presence of proteins P. For the homopolymer of A, only AA, AP and PP interactions are implemented.

| $ij$ | $\varepsilon_{ij} (k_B T)$ | $r_{c,ij} (\sigma)$ |
| --- | --- | --- |
| AA | 1.0 | 1.12 |
| BB | 1.0 | 1.12 |
| AB | 1.0 | 1.12 |
| AP | 1.5 | 2.5 |
| BP | 2.0 | 2.5 |
| PP | $\varepsilon_{PP} = 0.1 \text{ to } 2.0$ | 2.5 |

**TABLE II.**
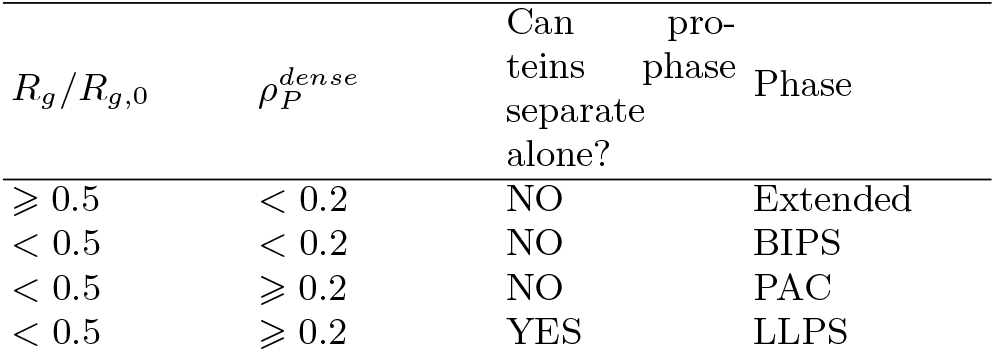
Criteria for determining the different phases of homopolymeric chromatin and protein system.

In our computational study, the simulated chromatin and proteins are confined within a cubic box with dimensions *L*_*x*_ = *L*_*y*_ = *L*_*z*_ = 100 *σ*, a size chosen to ensure that the chromatin adopts an extended configuration initially in the absence of proteins, as demonstrated in Fig.S1. The radius of gyration of the model chromatin, as defined below, is measured to estimate the required simulation box length:

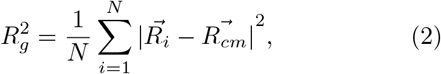

where 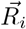 is the position vector of the *i*th monomer, and 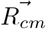 is the position vector of the chromatin’s center of mass. The *R*_*g*_ of the chromatin without proteins, *R*_*g*,0_, is approximately 30 *σ* (Fig.S1), substantiating the choice of box length at 100 *σ* to comfortably exceed the chromatin’s end-to-end distance, thus mitigating boundary effects on the system’s intrinsic dynamics. In our simulations, the protein-protein interaction strength (*ε*_*P* *P*_ ) is parameterized to vary from 0.1 to 2 *k*_*B*_*T* , with the total number of proteins (*N*_*P*_ ) ranging from 500 to 6000. This variation yields a protein density (*ρ*_*P*_ ) spanning from 0.0005 to 0.006 *σ*^*−*3^. Notably, at *ε*_*P* *P*_ = 2, protein condensation in the absence of chromatin is observed when *ρ*_*P*_ *≥* 0.0045, as evidenced in Fig.S2. The careful selection of these parameters for *ε*_*P* *P*_ and *ρ*_*P*_ facilitates an exhaustive exploration of the dynamics spanning Bridging-Induced Phase Separation (BIPS), Polymer-Assisted Condensation (PAC), and Liquid-Liquid Phase Separation (LLPS) regimes. Additionally, interactions between both monomers and proteins with the confining cubic box walls are modeled through a Lennard-Jones potential, as detailed in the model presented by Chaudhuri et al.^55^. This interaction is critical for elucidating the behavior of chromatin and proteins under varied conditions in a confined environment, enabling a controlled investigation into the phase behavior and structural organization of the system under study.

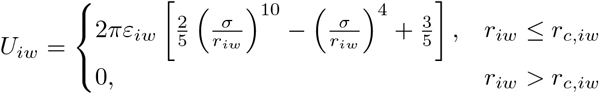

Here, *ε*_*iw*_ = 1*k*_*B*_*T* and *r*_*c*,*iw*_ = *σ*, where *i* stands for both monomers and proteins, *w* stands for the walls.

Our computational model is designed to be scale-free, affording the versatility to represent a polymer comprising 2000 monomers as DNA segments of variable lengths, ranging from 500 Kbp to 5 Mbp. Adjustments to all corresponding dimensions within the system are made proportionately. Selecting a DNA segment length of 500 Kbp necessitates setting the value of *σ* to correspond to 250 base pairs (bp) or equivalently, 85 nm. This scaling is pertinent given that, in eukaryotic systems, a single protein typically binds to an average DNA length of approximately 10 bp^56^, with the physical dimensions of proteins ranging between 3 to 6 nm^57^. Although our model does not precisely match the real-world resolution regarding the DNA segment length per protein binding, converting the protein density (*ρ*_*P*_ ) into concentration demonstrates that a *ρ*_*P*_ value of 0.002 *σ*^*−*3^ translates to a protein concentration of 5 nM. This concentration falls within the physiological range observed in cellular environments^57^. To model a DNA length of 500 Kbp alongside a more accurate representation of protein dimensions, a chromatin model consisting of 50,000 monomers would be necessary. However, achieving an extended chromatin configuration in the absence of proteins for such an expansive system would require increasing the simulation box size to at least 660 *σ*?this estimation arises from the scaling relationship *R*_*g*,0_ *∼ N* ^3*/*5^. Furthermore, to attain physiological protein concentrations, the inclusion of approximately 10^6^ proteins would be required. Simulations of this magnitude are computationally prohibitive; therefore, we have opted for a parameterization where *σ* equals 85 nm, enabling our simulations to cover a protein density range of 0.0005 *≤ρ*_*P*_ *≤*0.006.This corresponds to a protein concentration range of approximately 1.3 nM to 16 nM. By adopting this approach, we can conduct simulations that are computationally feasible and still yield valuable insights into biological phenomena within a constrained, yet relevant, parameter space..

### B. Methods

We perform molecular dynamics simulations in the NVT ensemble, integrating the equations of motion and maintaining constant temperature via a Langevin thermostat. Across all simulations, the Boltzmann constant times temperature (*k*_*B*_*T* ) is set to 1, with the friction coefficient (*γ*) also maintained at 1. The selected time step for our simulations is 0.01*τ* , defined as 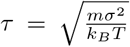 . For these simulations, we employ the Espresso-3.3.1 package^58^, and for visualization, we use the VMD package^59^. Each simulation initiates from an equilibrated chromatin configuration in the absence of proteins. Upon introducing proteins, we allow the system to equilibrate for 300, 000*τ* , followed by a production phase lasting 20, 000*τ* . To ascertain equilibrium, we monitor the radius of gyration (*R*_*g*_) of chromatin, as depicted in Fig.S3, with the equilibrium *R*_*g*_ value computed as an average over the production run. In cases with *ε*_*P* *P*_ = 0.1 and low protein densities (*ρ*_*P*_ *≤* 0.002), where the homopolymer size approximates that of the free polymer (*R*_*g*,0_) as observed in the simulation (refer to Fig.S3), we report both the mean *R*_*g*_ and its fluctuation throughout the simulation. For statistical robustness, averages are taken over 100 independent configurations after 300, 000*τ* , each spaced by 200*τ* , to calculate mean square spatial distances between monomers ( ⟨ *R*^2^(*s*) ⟩ ), mean contact probabilities ( ⟨ *P*_*c*_(*s*) ⟩ ), and generate contact maps. To determine the scaling exponents (*ν* and *α*), we fit the ⟨ *R*^2^(*s*) ⟩ and ⟨ *P*_*c*_(*s*) ⟩ to power-law expressions on a log-log scale, as illustrated in Figs.S4 and S5. All data analyses are performed using custom-developed C and Python programs.

## III. RESULTS

### A. Protein-protein interaction and protein concentration dictate chromatin organization

To dissect how *ε*_*P* *P*_ and *ρ*_*P*_ affect chromatin organization, first, we consider the scenario of a homogeneous chromatin. In particular, we systematically study the different organizations that can emerge as we tune *ε*_*P* *P*_ and *ρ*_*P*_ . Initially, we determine the equilibrium size of the chromatin by measuring the radius of gyration (*R*_*g*_). Fig.2(*a*) shows that the scaled radius of gyration (*R*_*g*_*/R*_*g*,0_) of the chromatin exhibits distinct behaviors at low and high *ε*_*P* *P*_ . For *ε*_*P* *P*_ *<* 1.5, protein-protein interactions are weaker than monomer-protein interactions, leading to chromatin collapse due to Bridging-Induced Phase Separation (BIPS) when *ρ*_*P*_ exceeds a certain threshold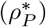 . This type of collapse has also been observed in the lattice-based SBS model, where proteins do not interact with each other^42^. As *ε*_*P* *P*_ increases, chromatin collapses for lower threshold 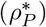 clearly indicating that protein-protein interaction has a significant effect on the BIPS regime. However, no swelling of the chromatin is observed in this regime. At *ε*_*P* *P*_ = 1.5, we observe a marginal swelling of the polymer for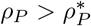 . Conversely, at *ε*_*P* *P*_ = 2, chromatin collapse is followed by a noticeable swelling phase when *ρ*_*P*_ *>* 0.002. Interestingly, in the absence of chromatin, a protein density range of 0.002 *< ρ*_*P*_ *<* 0.0045 is insufficient for proteins to form a condensate, as shown in Fig.,S2(*a*), indicating that this is outside the Liquid-Liquid Phase Separation (LLPS) regime. However, the presence of chromatin increases effective *ρ*_*P*_ , leading to droplet formation within the chromatin and causing it to swell, as depicted in Fig.,S2(*b*). This phenomenon is referred to as co-condensation of chromatin-protein or Polymer-Assisted Condensation (PAC) in the literature^36–40^. For *ρ*_*P*_ ) ⩾ 0.0045, proteins undergo LLPS, leading to a further increase in *R*_*g*_.

Although the scaled size of the chromatin in Fig. 2(*a*) illustrates the effect of protein density and proteinprotein interaction on the size of chromatin, it does not capture the influence of chromatin on protein organization.To address this, we compute the density of the condensed proteins, 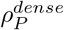 , defined as:

**FIG. 2.**
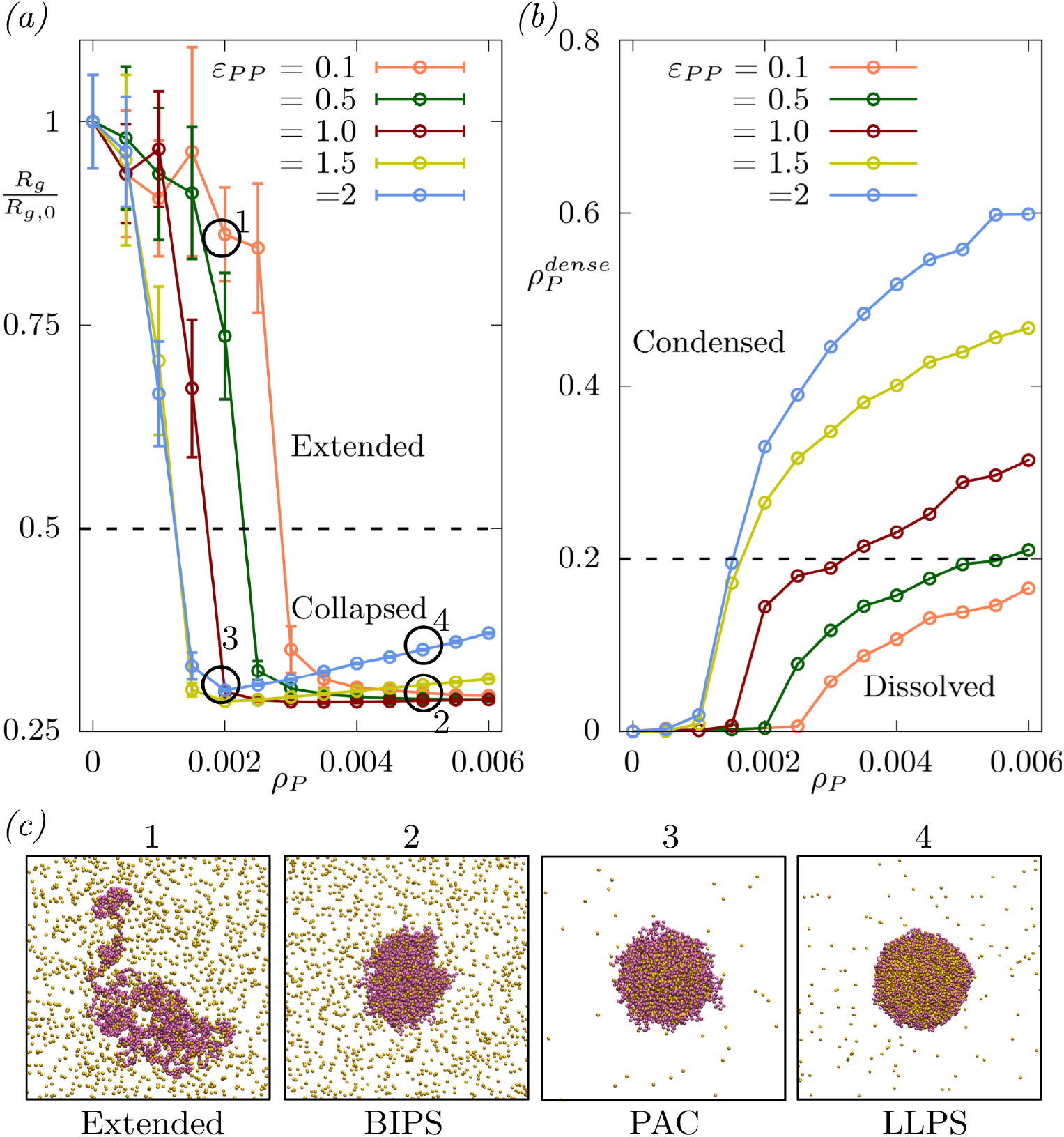
(*a*) Scaled radius of gyration (*R*_*g*_*/R*_*g*,0_) of the chromatin, and (*b*) density of the proteins inside the chromatin 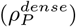, with increase in *ρ*_*P*_ at *ε*_*PP*_ =0.1 (orange), 0.5 (green), 1 (red), 1.5 (yellow), and 2 (blue). The dashed line denotes the chosen threshold for separating the phases of the chromatin and protein. (*c*) Configurations corresponding to the points marked as 1 (*ε*_*PP*_ =0.1, *ρ*_*P*_ =0.002), 2 (*ε*_*PP*_ =0.1, *ρ*_*P*_ =0.005), 3 (*ε*_*PP*_ =2, *ρ*_*P*_ =0.002), 4 (*ε*_*PP*_ =2, *ρ*_*P*_ =0.005) in (*a*).

**FIG. 3.**
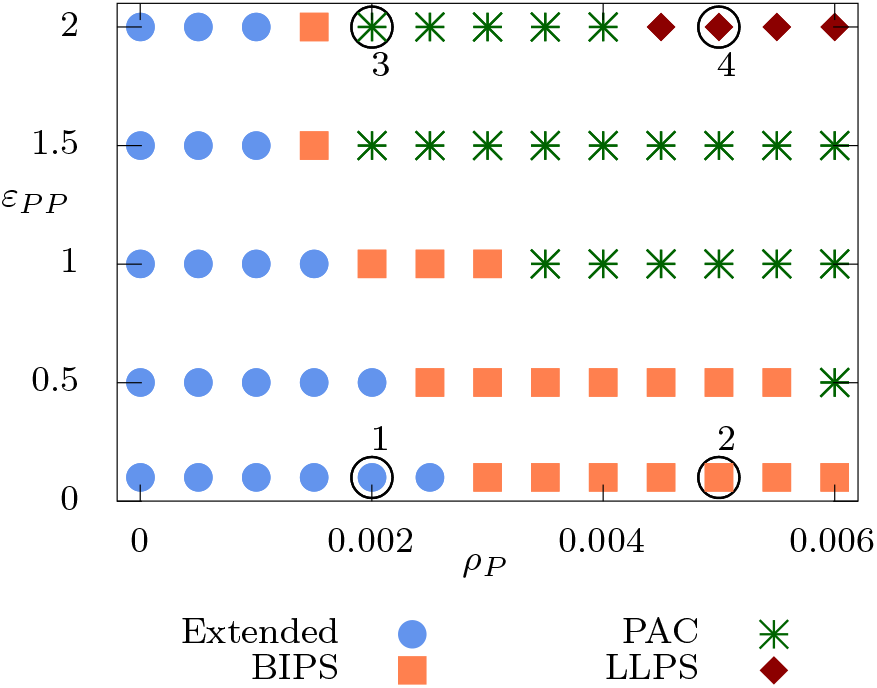
(*a*) Phase diagram of the homopolymer-protein system showing four distinct phases with variation in *ε*_*PP*_ and *ρ*_*P*_ . The encircled points marked as 1,2,3 and 4 (same as those shown in Fig. 2) are representatives of the four different phases.

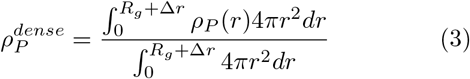

Here, *ρ*_*P*_ (*r*) denotes the protein density relative to the chromatin’s center of mass, and Δ_*r*_ represents the distance accounting for proteins adsorbed on the chromatin surface. We fix Δ_*r*_ at 2, and note that further increases do not qualitatively change the outcomes, as evidenced in Fig.S6. With an increase in *ρ*_*P*_ , the dense protein density 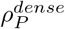 also increases; however, this rate of increase is notably more significant at higher *ε*_*P* *P*_ values, as illustrated in Fig.2(*b*). We establish a threshold of 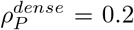 to distinguish between proteins in the ‘dissolved’ state and those in the ‘condensed’ state. This delineation is informed by the observation that, even at *ε*_*P* *P*_ = 0.1, a populated dilute phase of proteins remains at high *ρ*_*P*_ values, as shown in Fig.2(*c*), and the 0.2 threshold categorizes this phase of proteins as ‘dissolved.’ This differentiation is crucial for distinguishing between the PAC phase and the BIPS phase, as presented in the phase diagram (Fig.3). Consequently, through the combined analysis of protein behavior both in the absence and presence of chromatin, the two metrics *R*_*g*_*/R*_*g*,0_ and 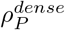 enable us to identify the extended, BIPS, PAC, and LLPS regimes within the system. We outline our criteria for determining the phase diagram in Table III A.

### B. *ε*_*P* *P*_ and *ρ*_*P*_ influence the genomic loci exponent (*ν*), contact probability exponent (*α*) and contact maps

While *R*_*g*_*/R*_*g*,0_ offers insight into the global size of chromatin, it does not elucidate how protein-protein interactions influence the distances between different chromatin segments. To probe this aspect, we assess the three-dimensional (3D) distance between chromatin segments as a function of genomic distance (⟨*R*^2^(*s*) *∼ s*^*ν*^ ⟩) and calculate the exponent (*ν*), as illustrated in Fig., S4. In Fig. 4, we examine the variation of the internal distance exponent (*ν*) with protein density (*ρ*_*P*_ ). At lower *ρ*_*P*_ values, an increase in *ε*_*P* *P*_ results in enhanced chromatin compaction, as indicated by a decrease in *ν*. Conversely, at higher *ρ*_*P*_ values, the differences between *ε*_*P* *P*_ = 0.1 and 2 become negligible. Throughout our simulations, *ν* spans from 0.33 to 0.6, which is consistent with the theoretical range for a self-avoiding polymer^60^ and closely parallels the experimentally observed chromatin range of 0.3–0.5^50^. The Strings and Binders Switch (SBS) model posits that *ν* can vary from 0 to 0.6^42^. To explore the lower bound of *ν*, we performed simulations with a homopolymer in the presence of purely repulsive proteins, finding that *ν* fluctuates between 0.4 to 0.6 (Fig.,S7). This range, higher than in scenarios with attractive *ε*_*P* *P*_ = 0.1, is attributed to protein-protein repulsion leading to chromatin expansion, suggesting that even protein volume exclusion results in a non-zero *ν* value. The observation of *ν* = 0 in the SBS model is ascribed to the lack of interaction between proteins. Nonetheless, the SBS model’s prediction of *ν* = 0 aligns with spatial distance measurements for the immunoglobulin heavy-chain locus, a segment of chromosome-14, in human pro-B cells, as reported by Jhunjhunwala and colleagues, who observed *ν* = 0.1, not *ν* = 0, and applied the Multi-Loop-Subcompartment (MLS) model to interpret their findings^51^. The MLS model envisages chromatin as comprising subcompartments (1-2 Mbp) interconnected by smaller loops (60-250 kbp) through variable-sized linkers (60-250 kbp), integrating aspects of both loop extrusion and phase separation models^61^. Barbieri and colleagues’ observation of *ν* = 0 is explained by the absence of volume exclusion, permitting multiple protein molecules to bind to the same chromatin site and form loops^42^. This indicates that a synthesis of loop formation and phase separation mechanisms could account for the instances of *ν <* 0.33 reported in some studies^51,62^. However, in the absence of loop-like structures and with volume exclusion between proteins, as in our model, *ν* = 0.33 represents the minimum achievable genomic loci exponent.

**FIG. 4.**
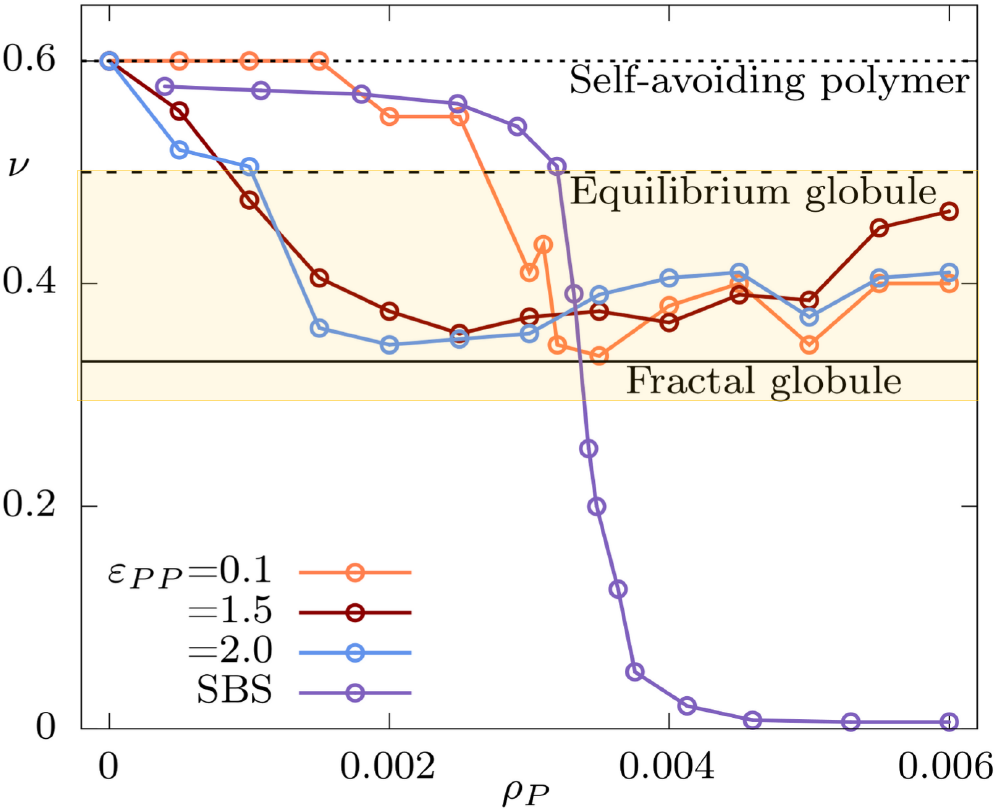
Genomic loci exponent (*ν*) of the homopolymeric chromatin with increase in *ρ*_*P*_ at *ε*_*PP*_ =0.1 (orange), 1.5 (red), and 2 (blue). The purple line is the value of *ν* extracted from Fig. 1 D of the SBS model^42^. The shaded region shows the range of *ν* reported in most of the experiments^50^.

**FIG. 5.**
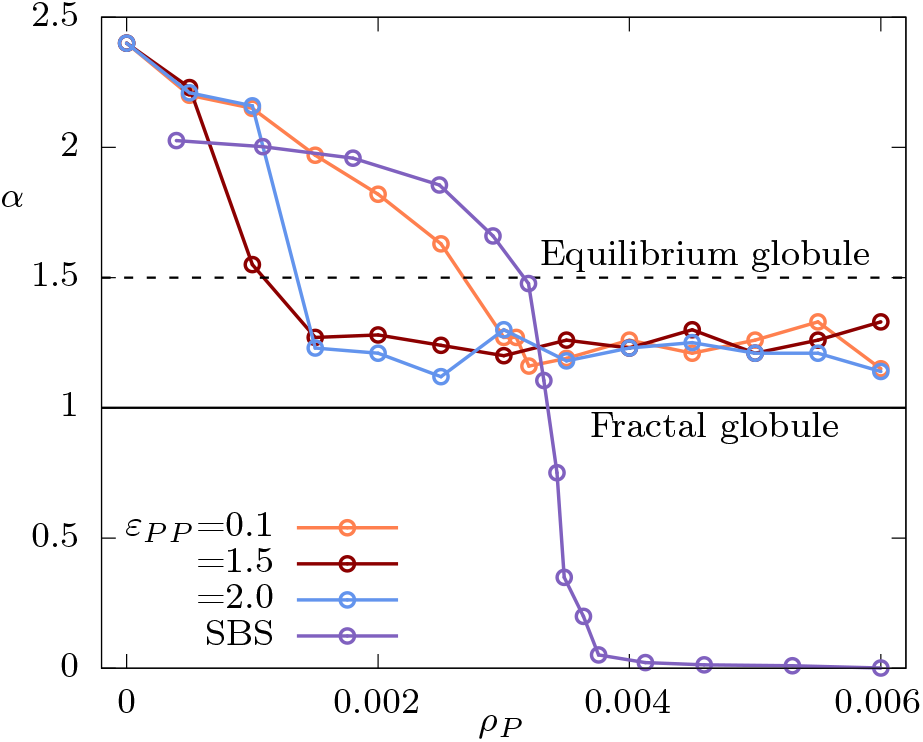
Contact probability exponent (*α* ) of the homopolymeric chromatin with increase in *ρ*_*P*_ at *ε*_*PP*_ =0.1 (orange), 1.5 (red), and 2 (blue). The purple line is the value of *α* extracted from Fig. 1 F of the SBS model^42^.

Similar to the trends observed with the internal distance exponent (*ν*), the contact probability exponent (*α*) also exhibits a more pronounced decrease with increasing protein density (*ρ*_*P*_ ) when the protein-protein interaction energy is set at *ε*_*P* *P*_ = 2, compared to *ε*_*P* *P*_ = 0.1, especially for *ρ*_*P*_ *≤*0.002 (as demonstrated in Fig.,S5(*a*)). However, discerning the impact of *ε*_*P* *P*_ on *α* becomes more complex for *ρ*_*P*_ *≥* 0.002. Yet, when examining the absolute value of the contact probability *P*_*c*_(*s*), a clear decline in contact probability is observed as *ε*_*P* *P*_ increases from 0.1 to 2, as illustrated in Fig.,S5(*b*). This difference becomes even more pronounced when analyzing the contact maps in Fig.6. At *ε*_*P* *P*_ = 0.1, elevating *ρ*_*P*_ from 0.002 to 0.005 enhances bridging, leading to increased contacts between monomers, as shown in Fig.6(*a*) and (*b*). In contrast, at *ε*_*P* *P*_ = 2, a similar increase in *ρ*_*P*_ results in protein droplet formation, which subsequently diminishes monomer contacts, as evident in Fig.6(*c*) and (*d*)). The configurations depicted in Fig.2(*c*), along with those in Fig., S8, support this interpretation. Our analysis of contact maps offers an indirect method for inferring the competition between protein-protein interactions and protein-DNA interactions from Hi-C maps. If a knockdown assay shows a reduction in contact maps, it indicates that the particular protein may be acting as a bridger, i.e., a protein with higher DNA binding affinity than self-association affinity. Conversely, if the deletion of a protein leads to an increase in contact intensities, it is likely that the deleted protein was suppressing contacts between monomers by forming stronger associations with itself. For instance, the knockdown of the HP1 variant found in Drosophila (HP1a) results in an increase in contacts in the Hi-C map^63^, suggesting that Hp1a-Hp1a interaction is stronger than Hp1a-DNA interaction. This interpretation aligns with in vivo experiments by Strom and colleagues, who observed that HP1a protein forms liquid droplets in early Drosophila embryos^64^, further illustrating the complex interplay between protein associations and chromatin structure.

**FIG. 6.**
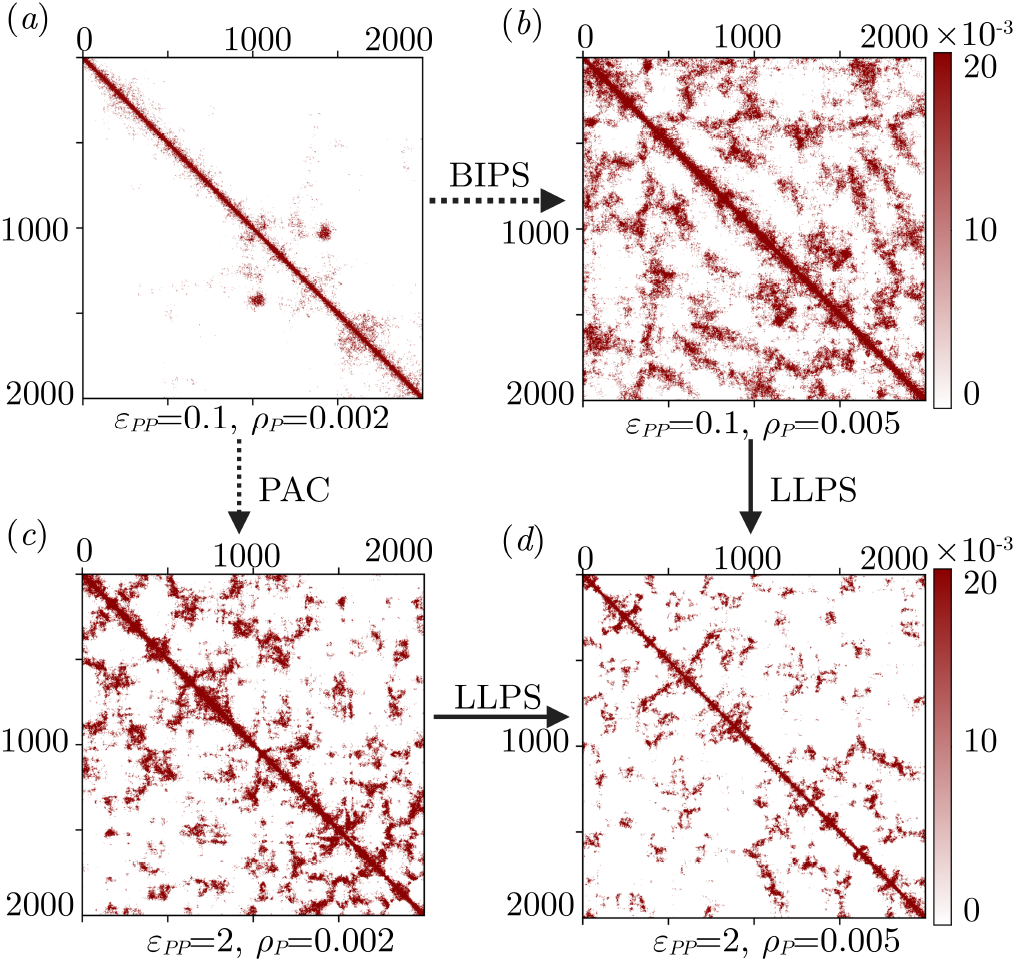
Contact maps between the monomers corresponding to the configurations shown in Fig. 2(*c*).

### C. Sequence heterogeneity affects the internal organization of chromatin

Next, we focus on the impact of sequence heterogeneity by analyzing three chromatin models as depicted in Fig.1, maintaining *ε*_*P* *P*_ at 0.1 across all sequences while varying *ρ*_*P*_ . In the block copolymer model, we substitute half of the A monomers (specifically, those in the index range 1000 to 2000) with B monomers. As detailed in Sec.II, B monomers exhibit a stronger affinity for proteins compared to A monomers. This differential affinity results in an elevated average monomer-protein interaction energy for the block copolymer relative to the homopolymer. Consequently, the block copolymer undergoes collapse at a lower protein density (*ρ*_*P*_ = 0.002) compared to the threshold density for the homopolymer collapse (as illustrated in Fig.7(*a*)). However, for *ρ*_*P*_ *>* 0.004, the scaled radius of gyration (*R*_*g*_*/R*_*g*,0_) for both polymers becomes virtually indistinguishable. While the homopolymer and block copolymer differ in average monomer-protein interaction energy, the alternative polymer shares the same average interaction energy as the block copolymer but exhibits distinct sequence heterogeneity. Fig. 7(*a*) indicates that the sizes of the block copolymer and the alternative polymer are nearly identical across the protein density spectrum. Yet, a detailed examination reveals that the alternative polymer undergoes collapse in a switch-like fashion, reminiscent of the homopolymer, whereas the block copolymer demonstrates a more gradual size reduction. At *ρ*_*P*_ = 0.001, the block copolymer sequence displays a higher *R*_*g*_*/R*_*g*,0_ (0.55) compared to the alternative polymer (0.31), suggesting that half of the block copolymer may be in a collapsed state while the other half remains extended. This observation leads to speculation that despite identical average monomer-protein interaction energies, sequence heterogeneity significantly influences the local structure and compaction dynamics of chromatin. The differential behavior between the block and alternative polymers underscores the importance of monomer sequence distribution in dictating chromatin folding and compaction.

**FIG. 7.**
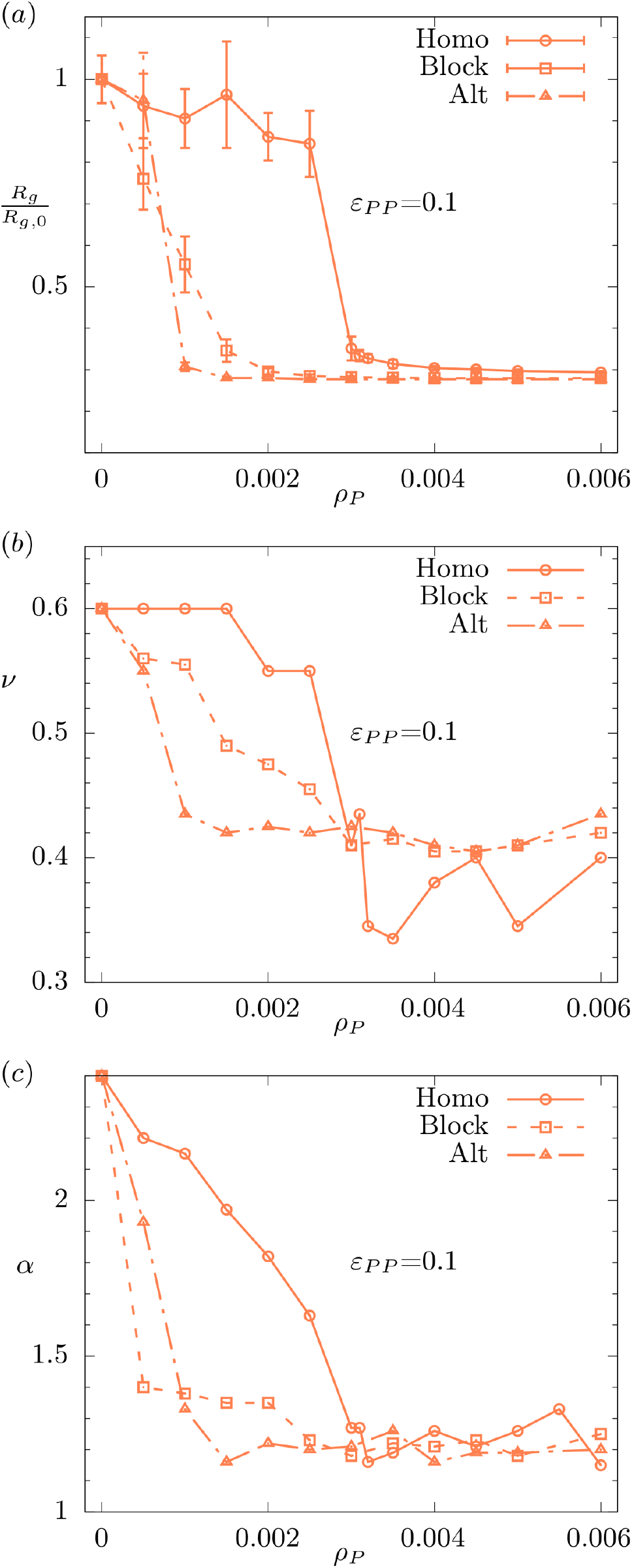
(*α*) Scaled radius of gyration (*R*_*g*_*/R*_*g*,0_), (*b*) genomic loci exponent (*ν*), and (*c*) Contact probability exponent (*α*) as a function of *ρ*_*P*_ for homopolymer (solid line), block copolymer (dashed line) and alternative polymer (dash-dotted line). *ε*_*PP*_ =0.1 for all cases.

To validate our hypothesis regarding the distinct local structures of different chromatin sequences, we examine the genomic loci exponent (*ν*) and the contact probability exponent (*α*) as presented in Fig. 7(*b*) and (*c*). At *ρ*_*P*_ *<* 0.003, the disparity in *ν* across the three sequences reflects the variation observed in their respective *R*_*g*_ values, substantiating our conjecture that certain regions within the block copolymer exhibit more expansion than others. A qualitatively similar trend is noted in *α* as well. Nonetheless, a discrepancy emerges between *ν* and *α* at *ρ*_*P*_ *<* 0.001. In this domain, although *ν* is elevated for the block copolymer, *α* is higher for the alternative polymer. This divergence might stem from the more pronounced fluctuations in chromatin structure at very low protein densities, nuances that are not adequately captured when deriving the exponents *ν* and *α* from the mean square spatial distances between monomers ( ⟨ *R*^2^(*s*) ⟩) and mean contact probabilities ( ⟨ *P*_*c*_(*s*) ⟩ ). Crucially, at elevated *ρ*_*P*_ levels, no discernible distinction is observed between the block copolymer and alternative polymer sequences, even when examining local metrics such as *ν* and *α*. This outcome suggests that, despite initial differences in local structure and compaction dynamics at lower protein densities, the eventual structural configurations of both sequences converge at higher *ρ*_*P*_ , indicating a uniformity in chromatin compaction behavior under high protein concentration conditions.

To elucidate the sequence-specific characteristics of different chromatin models at high protein density, we analyze their contact maps. Figures 8(*a*) and (*b*) illustrate that, whereas the homopolymer exhibits uniform contact frequencies, the block copolymer displays increased contact frequencies within the segment containing B monomers and reduced frequencies in the segment with A monomers, as evidenced by the contrast between the dashed and solid boundary regions. Conversely, the alternative polymer demonstrates more uniform contact frequencies, akin to those observed in the homopolymer’s contact map (Fig.8(*c*)). A detailed examination of the contact maps, particularly between monomers 1000 to 2000 (Fig.8(*d*)–(*f* )), uncovers notable differences between the sequences. The block copolymer sequence exhibits an increase in contact frequencies among off-diagonal indices compared to the homopolymer, whereas the alternative sequence shows a decrease in contact frequencies between off-diagonal indices relative to the homopolymer. This variation in off-diagonal contact frequencies between the block copolymer and alternative polymer underscores a divergence in their internal organization. The enhanced off-diagonal contacts in the block copolymer suggest a propensity for interactions between distinct regions, possibly reflecting a more segmented or compartmentalized internal structure. On the other hand, the reduced off-diagonal contacts in the alternative polymer indicate a more homogenous internal distribution, mirroring the homopolymer’s behavior.

**FIG. 8.**
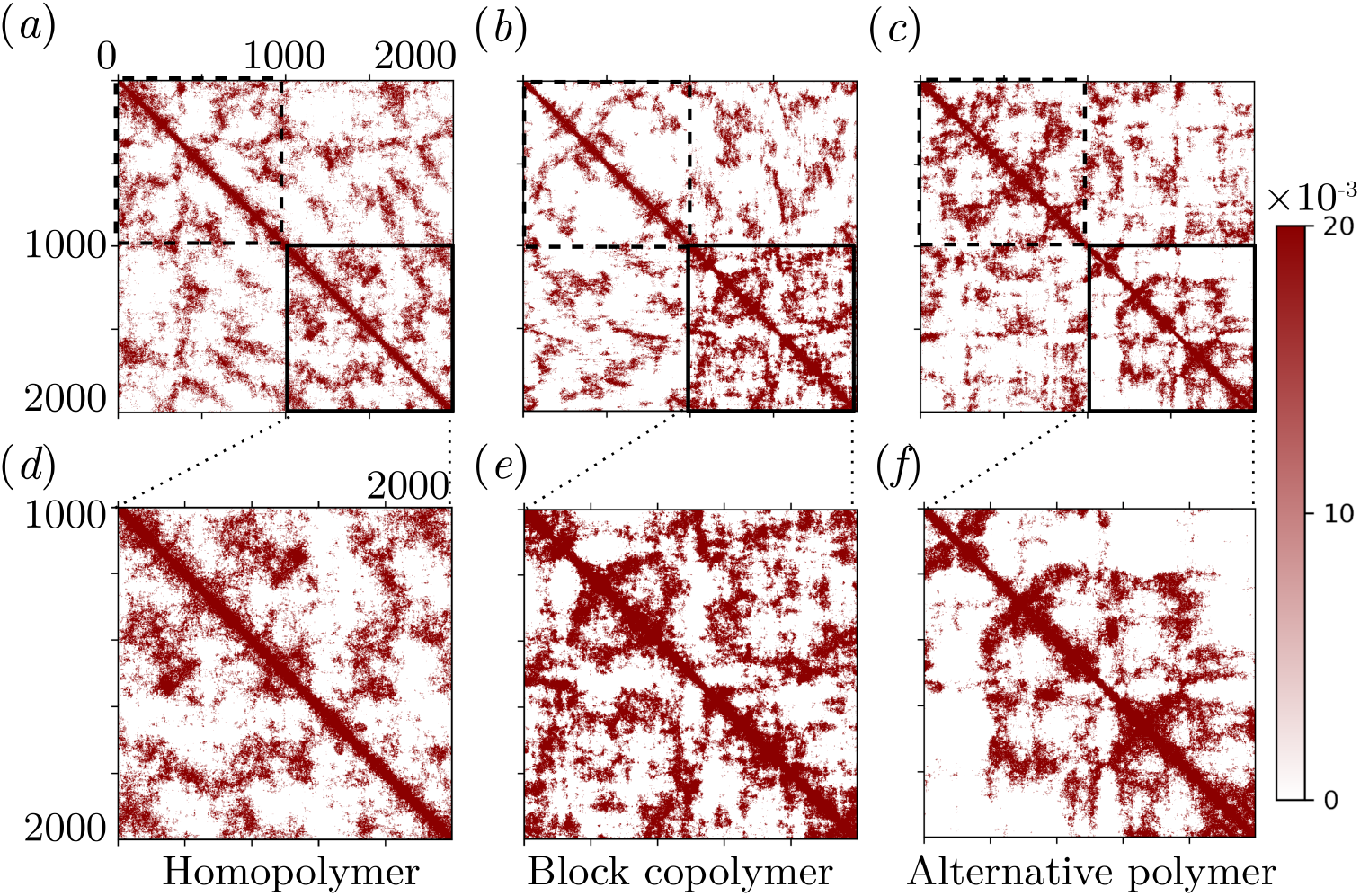
Contact maps between the monomers at *ε*_*PP*_ =0.1 and *ρ*_*P*_ = 0.005 for (*a*) homopolymer and (*b*) block copolymer (*c*) alternative polymer. In (*d*)–(*f* ), we show the enlarged view of the contact maps between 1000-2000 monomer indices in (*a*)–(*c*).

To gain further insight into the organization of monomers A and B within the polymers, we investigate the density of these monomers (*ρ*(*r*)) in relation to the center of mass of the polymer. Figures 9(*a*) and (*b*) illustrate that the block copolymer demonstrates predominantly distinct regions of A and B monomers, leading to a layered organization throughout the polymer (refer to Fig.,S9 for additional detail). In contrast, the alternative polymer shows overlapping densities for A and B monomers, as observed in Figures 9(*c*) and (*d*)), resulting in a more uniform structural composition. The underlying reason for this discrepancy in organizational structure can be elucidated through both thermodynamic and kinetic considerations. From a thermodynamic viewpoint, situating the B monomers at the core of the block copolymer minimizes the interfacial tension between different types of monomers. On the kinetic front, the B monomers, which exhibit a stronger attraction to proteins, tend to collapse initially, subsequently drawing in the A monomers, which possess a weaker affinity for proteins. This sequential compaction fosters a layered architecture within the block copolymer, as visualized in Fig.S10. Such differences in monomer organization and density distribution highlight the nuanced interplay between monomer composition, protein binding affinities, and polymer structure. The block copolymer’s layered configuration is indicative of a spatially heterogeneous structure, where monomer type directly influences local compaction and organization. Conversely, the more homogeneous structure of the alternative polymer suggests a more uniform distribution of protein interactions across the polymer, irrespective of monomer type. These observations underscore the significance of sequence arrangement and protein affinity in determining the overall architecture, spatial organization, and functionality within the chromatin models, underlining the impact of sequence heterogeneity on chromatin architecture at a molecular level.

**FIG. 9.**
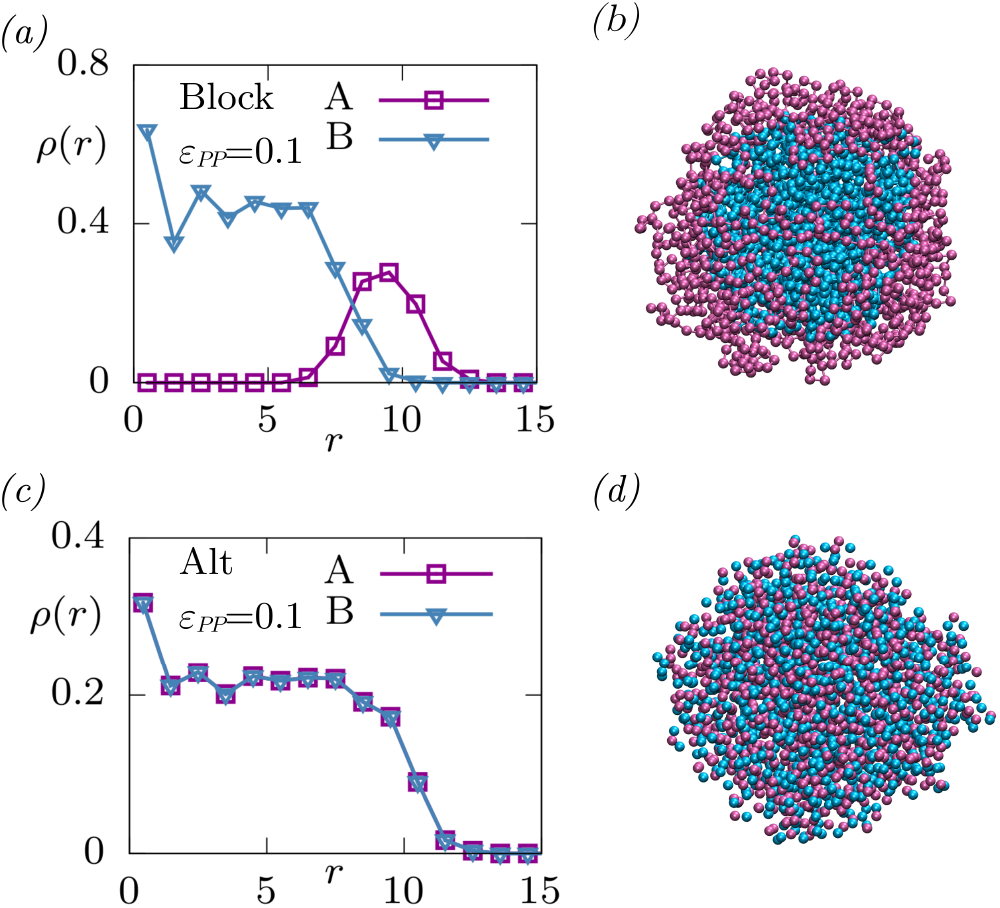
Density of the monomer A (magenta □), monomer B (cyan ▽) from the center of mass of the polymer for (*a*) block copolymer and (*b*) alternative polymer. In (*c*) and (*d*) we show the configurations of the polymers in (*a*) and (*b*). *ε*_*PP*_ =0.1 and *ρ*_*P*_ =0.005 for both the cases. Configurations including proteins are shown in Fig. S9.

## IV. DISCUSSION

In this paper, we explore the role of chromatin sequence and the interaction between proteins in shaping its organization. Using simple polymer-based models, we uncover a novel mechanism for chromatin self-organization by exclusively adjusting protein-protein interactions. Our findings suggest that tuning proteinprotein interactions alone is sufficient to drive distinct self-organization patterns based on polymer sequence. Additionally, the chromatin serves as a nucleating site, facilitating protein condensation at lower densities than those required for the bulk phase separation of proteins alone. Furthermore, exponents describing contact probabilities along the chromatin are strongly influenced by both sequence and protein-protein attraction, potentially leading to a reinterpretation of Hi-C maps. Phase diagrams of structural and dynamical exponents, as a function of protein-protein interaction strength and density, unveil the complex interplay between sequence and intermolecular interactions in chromatin organization and phase separation. Transitioning to chromatin, a heteropolymer with varied interacting sites, its spatial organization relative to interacting proteins garners significant interest. Specific and non-specific interactions between proteins and chromatin domains in higher-order interphase chromatin organization are implicated in numerous functions and disease pathologies. Despite insights from Hi-C maps and simple polymer models, the role of binding protein-chromatin interactions in organizing chromatin hetero domains remains understudied. In the context of binding protein-chromatin interactions, it is noteworthy to highlight the recent investigation conducted by Adame-Arana and colleagues^24^. They explored the microphase separation of chromatin in the presence of a binding protein that binds reversibly to the active region of the chromatin. However, it is important to note that their model did not incorporate interactions among the proteins. Soft interactions among binding proteins could enhance interactions with different chromatin parts, promoting a hierarchical organization of chromatin folding. Microphase separation of block copolymers serves as a model for understanding disordered proteins, RNA segment localization, chromosome structuring, and euchromatin-heterochromatin compartmentalization.

We employed a homopolymer as a model system to investigate the mechanisms of Bridging-Induced Phase Separation (BIPS), Polymer-Assisted Condensation (PAC), and Liquid-Liquid Phase Separation (LLPS) within a system comprising chromatin and interacting proteins. Through measurements of the radius of gyration (*R*_*g*_), distinct chromatin behaviors were observed under varying protein-protein interaction energies (*ε*_*P* *P*_ ) and protein densities (*ρ*_*P*_ ). At lower *ε*_*P* *P*_ values, weaker protein-protein interactions led to chromatin collapse due to BIPS beyond a specific *ρ*_*P*_ threshold. This collapse mechanism at very low *ε*_*P* *P*_ values is similar to the finding from the lattice-based Strings and Binders Switch (SBS) model, which does not account for protein-protein interactions. However, as *ε*_*P* *P*_ increased, a more rapid chromatin collapse was noted, along with a shift in the collapse threshold to lower *ρ*_*P*_ , and significant swelling of the chromatin at higher *ε*_*P* *P*_ values, indicative of PAC phenomena.To gain further insight into chromatin’s influence on protein organization, we analyzed the density of condensed proteins. This analysis revealed that increased protein density enhances dense protein density 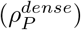, especially at higher *ε*_*P* *P*_ values. The analysis helped distinguish between the dissolved and condensed states of proteins. In conjunction with chromatin behavior in the absence of proteins, it allowed us to delineate extended, BIPS, PAC, and LLPS regimes within the system. Our findings contrast with those reported by Sommer and colleagues, mainly due to differences in fixed interaction strengths and the observed collapse-swelling transitions in chromatin. Investigations into the genomic loci exponent (*ν*) and the contact probability exponent (*α*) further elucidated internal structural differences among the chromatin models, with notable distinctions in contact frequencies and monomer densities.

Our phase diagram demonstrates a notable deviation from that reported by Sommer and colleagues^40^ in a key aspect. We maintain a constant monomer-protein interaction strength at 1.5 *k*_*B*_*T* and modulate the proteinprotein interaction strength along with protein density. Conversely, Sommer and colleagues^40^ fix the proteinprotein interaction strength at 1.1 *k*_*B*_*T* and vary the monomer-protein interaction strength as well as protein density. Despite these differences, Sommer and colleagues similarly report a collapse-swelling transition in the radius of gyration (*R*_*g*_) as protein density increases. Notably, at a monomer-protein interaction strength of 1 *k*_*B*_*T* , they identify a high-protein (HP) minority dense phase situated between the ‘no droplet’ phase and the ‘droplet’ phase. We correlate their ‘no droplet’ phase, ‘HP minority’ phase, and ‘droplet’ phase with the extended, Bridging-Induced Phase Separation (BIPS), and Polymer-Assisted Condensation (PAC) phases, respectively, in Fig. 3. However, their depiction of the droplet phase includes a significantly populated dilute phase, a variance that may be attributed to the protein density range (0 to 0.06) they simulated, which is an order of magnitude higher than our range (0 to 0.006). Both investigations probe only a fragment of the comprehensive three-dimensional phase diagram that incorporates monomer-protein interaction, protein-protein interaction, and protein concentration. The biological implications of these studies, particularly in relation to the HP1 protein, warrant further examination. Variations in monomer-protein interaction could indicate changes in histone methylation marks on heterochromatin where HP1 binds, while alterations in protein-protein interaction may reflect phosphorylation modifications on the HP1 protein. Specifically, our findings offer insights into in vitro experiments involving the HP1*α* and *λ*-DNA system as studied by Narlikar and colleagues^65,66^. Wild-type HP1*α* protein undergoes droplet formation at concentrations ranging from 400 to 800 *μ*M, depending on the salt concentration (40 mM to 75 mM KCl) in the absence of DNA. In contrast, Larson and colleagues observed that an N-terminal phosphorylated mutant of HP1*α* (nPhos-HP1*α*) forms droplets at a concentration of 60 *μ*M in the absence of DNA, indicating heightened protein-protein interaction for nPhos-HP1*α* compared to the wild-type. Furthermore, single-tethered experiments revealed that wild-type HP1*α* compacts DNA at a concentration of 1 *μ*M, whereas nPhos-HP1*α* requires 50 *μ*M for similar compaction. Additionally, they noted the compaction rate of nPhos-HP1*α* at 1 *μ*M/s, in contrast to 2.25 *μ*M/s for wild-type HP1*α*. These findings imply that the monomer-protein interaction is more pronounced for wild-type HP1*α* than for nPhos-HP1*α*. Consequently, we speculate that the wild-type HP1*α* and *λ*-DNA system predominantly reside within the BIPS regime, while the nPhos-HP1*α* and *λ*-DNA system largely fall within the PAC or co-condensation regime.

Here, we discuss the work by Tortola and colleagues, who investigated the role of protein-protein interaction and protein-DNA interaction in the context of heterochromatin and the HP1 protein^67^. The authors utilized a lattice gas model to study the role of HP1-H3K9me3 and HP1-HP1 interactions, aiming to understand the thermodynamics and kinetics of heterochromatin droplet formation. They varied HP1-HP1 interaction from 1 to 3 *kJ/mol* (equivalent to 0.4 to 1.2 *k*_*B*_*T* ) and HP1-H3K9me3 interaction from 0.1 to 0.6 *kJ/mol* (equivalent to 0.04 to 0.24 *k*_*B*_*T* ). In alignment with our findings and those of Sommer and colleagues^40^, they reported a reduction in *c*_*sat*_ in the presence of DNA. The authors also observed that the coil-globule transition of chromatin is consistent with both the BIPS and PAC mechanisms by simulating a model with HP1-HP1 interaction set to zero and HP1-H3k9me3 set to 1 *kJ/mol*. This corresponds well with our findings in Fig.2(*a*). They further reported a dependence of dilute phase protein density on the total density of proteins, as expected for a multicomponent system. Our dependence of dense phase protein density (Fig.2(*b*)) is consistent with this result. Despite these agreements, our interpretation of HP1a and HP1*α* proteins differs. The authors report that HP1a-HP1a interaction strength (1.8 kJ/mol) is weaker, corresponding to the HP1*α*-Hp1*α* interaction strength (2.6 kJ/mol). These interaction strengths are derived from their analysis of *c*_*sat*_ from turbidity assay experiments^68^, and mapping the *c*_*sat*_ to their phase diagram obtained from simulations. However, the authors report a *c*_*sat*_ of 7 *μ*M for HP1*α*, while the *c*_*sat*_ for HP1*α* in the absence of DNA is 400-800 *μ*M^66^. Our interpretation suggests that HP1*α*-DNA interaction is stronger than HP1*α*-HP1*α* interaction (BIPS), and HP1a-DNA interaction is weaker than HP1a-HP1a interaction (PAC, also see Sec. III B).

Furthermore, we investigated sequence heterogeneity using both block and alternative polymer models, reveal that the arrangement of sequences significantly influences the internal organization and compaction of chromatin. The block copolymer, featuring distinct regions of A and B monomers, exhibited a layered organization, while the alternative polymer displayed a more uniform structure due to overlapping densities of A and B monomers. The A and B monomers are different only from the perspective of the binding proteins and this organization of block copolymers in a corona-like architecture is primarily driven by the proteins interacting with A and B in a differential manner. The heteropolymers (both block and alternative) undergoes collapse at much lower protein density compared to homopolymer and is again driven by the proteins binding to A and B differently. For lower protein density, the contact probability exponents is significantly different for homo and heteropolymers and this can have significant impact on interpreting the Hi-C data. This difference arises from a combination of thermodynamic and kinetic factors, such as reduced interfacial tension, and kinetic factors, including the sequential compaction of monomers with varying affinities for proteins. These findings emphasize the crucial role of monomer sequence distribution and protein binding affinities in determining chromatin folding, compaction, and overall architectural organization. They underscore the impact of sequence heterogeneity on chromatin architecture at the molecular level.

In conclusion, our investigation presents a targeted and effective strategy for influencing chromatin organization by manipulating protein-protein interactions. The comparisons with equilibrium polymer models further highlight the significance of heterogeneity in our model, where the sequence of monomers with distinct affinities for proteins results in non-uniform compaction and distinct organizational patterns. Overall, our study not only addresses important questions in chromatin organization but also offers insights into the role of protein-protein interactions in complex biological systems and how tuning of the same can significantly impact the conforma.

## Acknowledgments

S.C. is supported by the Ramalingaswami Re-entry Fellowship (BT/HRD/35/02/2006), a re-entry scheme of the Department of Biotechnology, Ministry of Science and Technology, Government of India, and Department of Atomic Energy (DAE), Government of India, via Apex project to The Institute of Mathematical Sciences (IMSc), Chennai. The simulations were carried out on the supercomputing machines Annapurna and Nandadevi at the Institute of Mathematical Sciences.

