## Supplementary material for "Role of protein-protein interactions on model chromatin organization": supp_info

### Supporting information: Role of protein-protein interactions on model chromatin organization

(Dated: 31 January 2024)

[a)]Electronic mail:

[b)]Electronic mail:

[c)]Electronic mail:

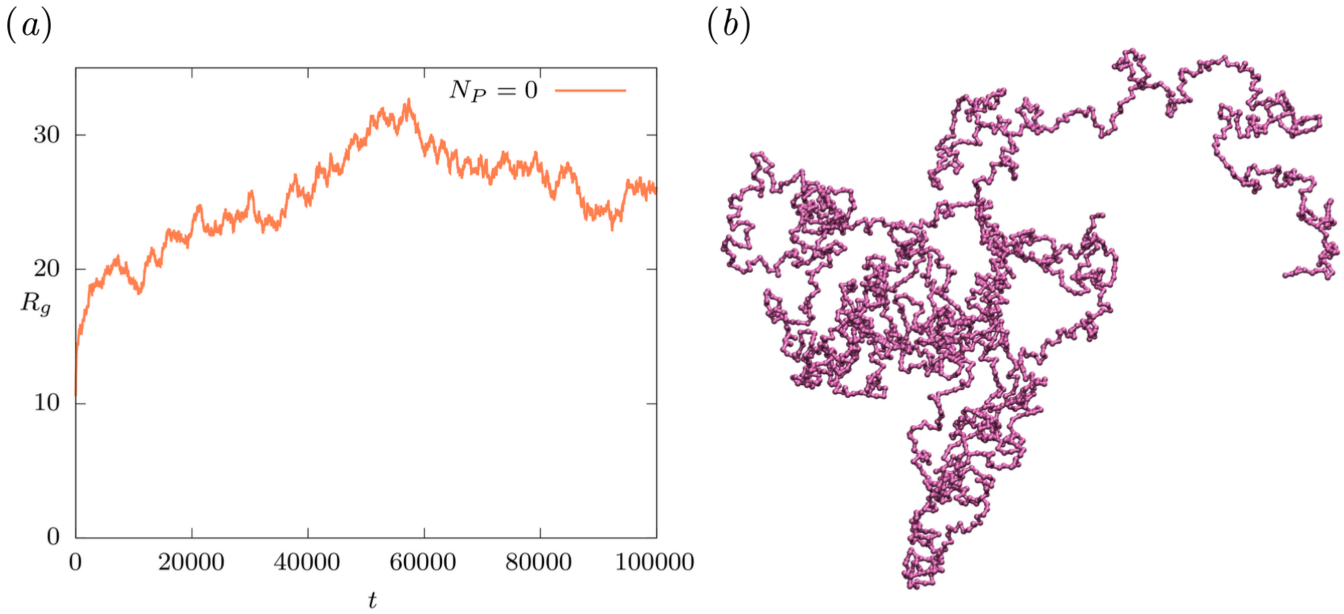


Fig. S1. $(a)$ Radius of gyration $(R_g)$, and $(b)$ configuration of the polymer in the absence of proteins.

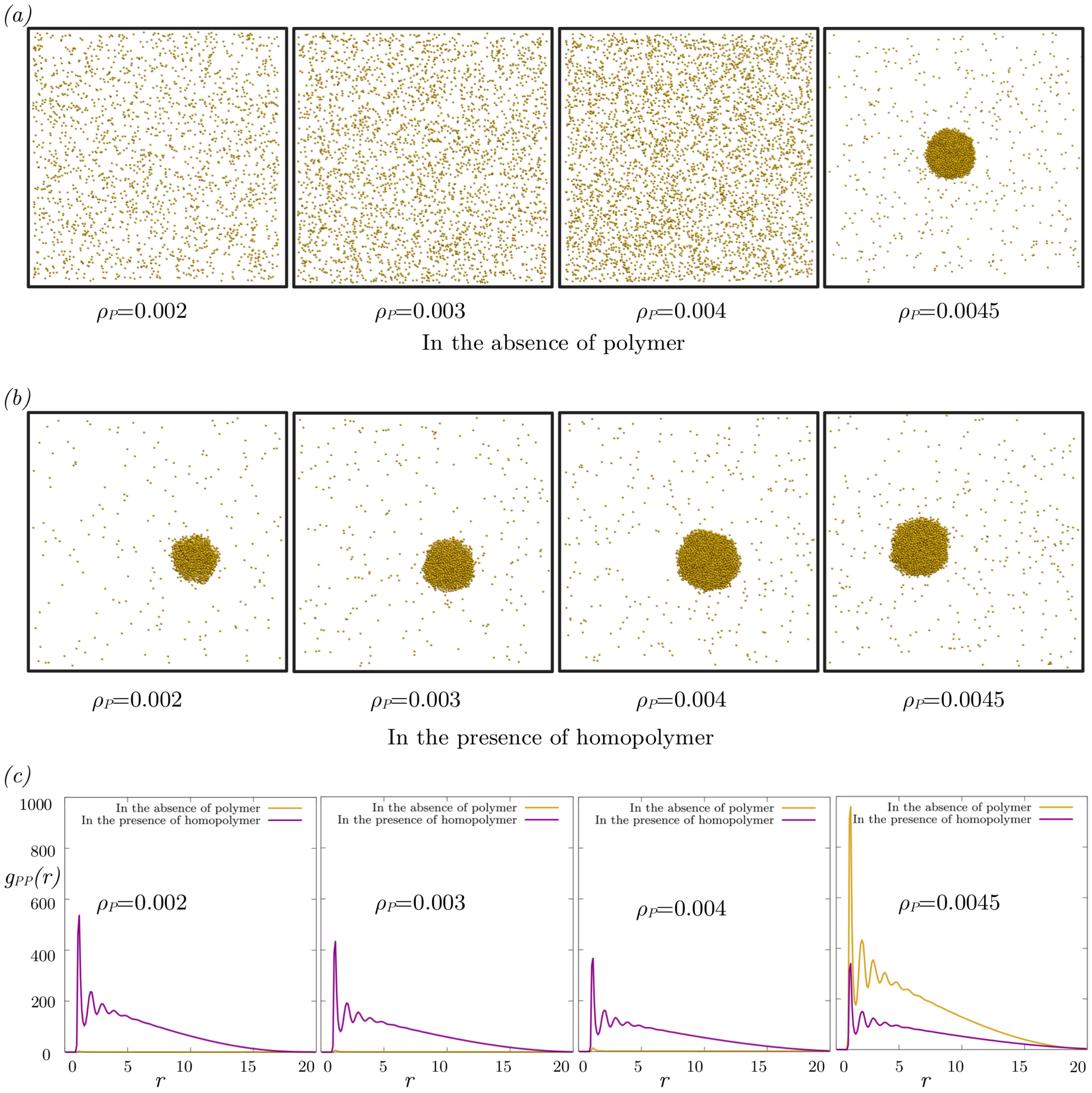


Fig. S2. Configurations of binders $(a)$ in the absence of polymer and $(b)$ in the prsence of homopolymer. $(c)$ radial distibution function between the binders $(g_{PP}(r))$ in the absence of polymer (yellow), and in the presence of homopolymer (magenta) for different $\rho_P$ values. $\varepsilon_{PP}$=2 for all the cases.

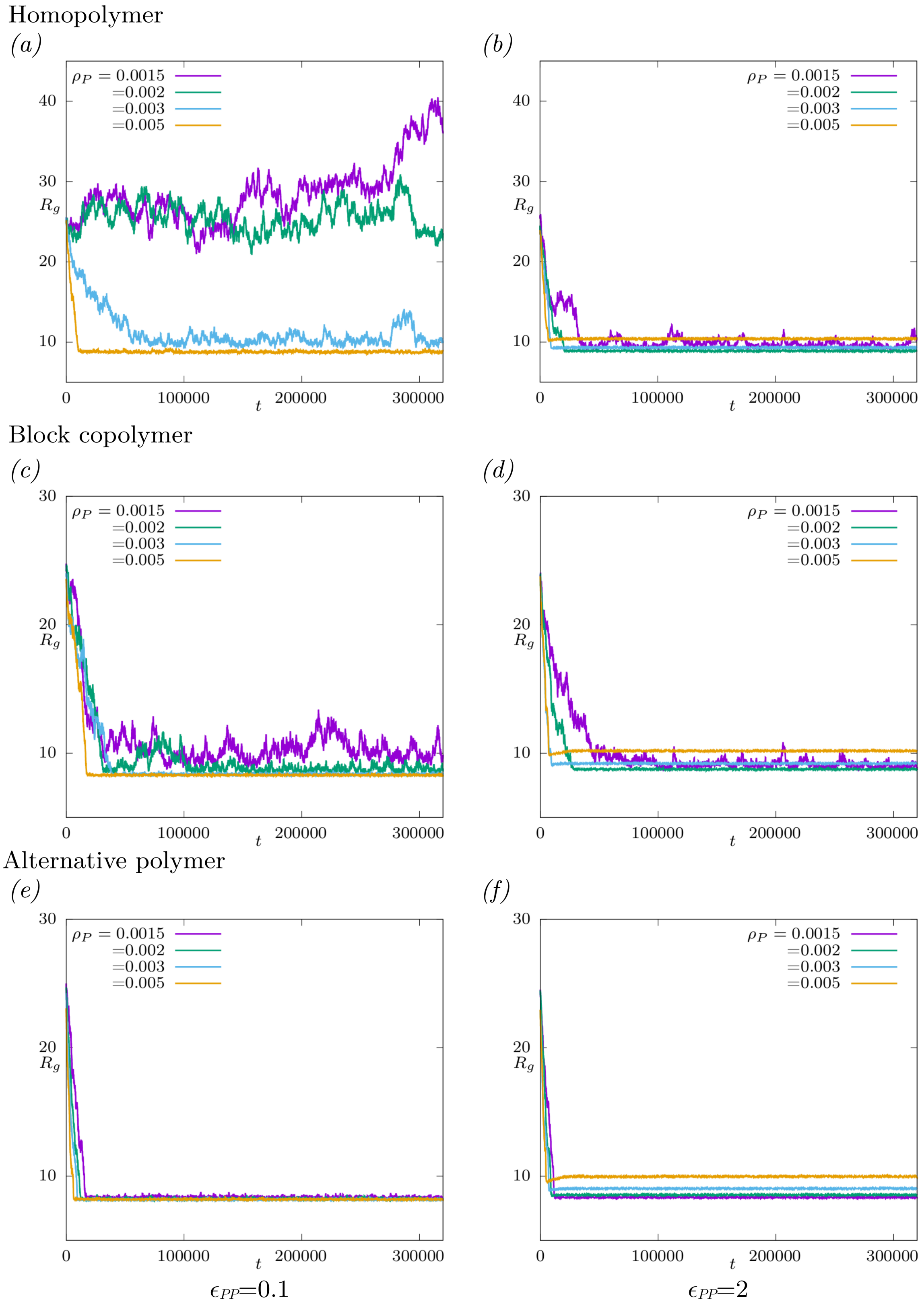


Fig. S3. Equilibriation of $R_g$ for homopolymer ($a$–$b$), block copolymer ($c$–$d$), and alternative polymer ($e$–$f$) at different values of $\rho_P$ and $\varepsilon_{PP}$.

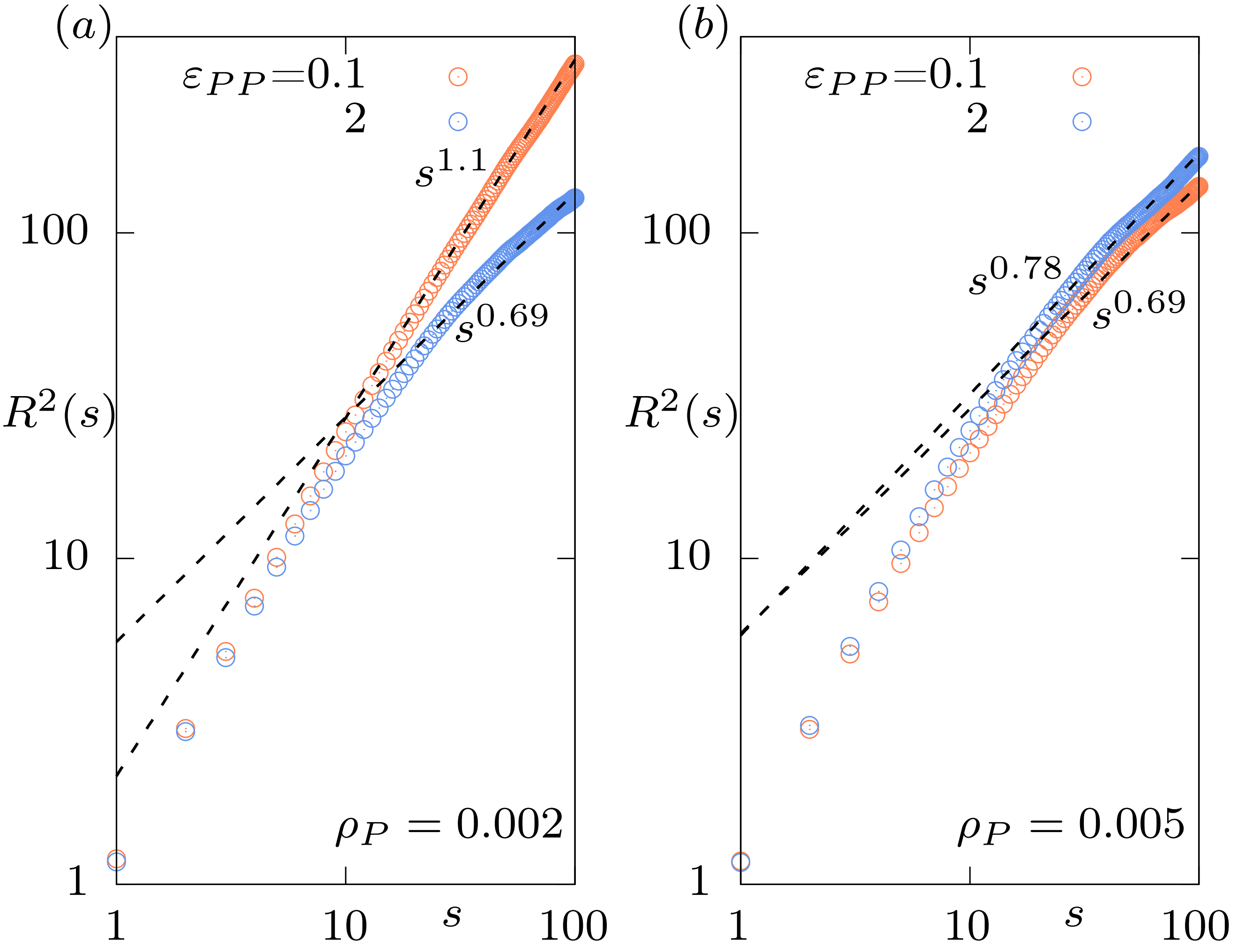


Fig. S4. Effect of $\varepsilon_{PP}$ on the inter-monomer distance of a homopolymer: $R^2(s)$ as a function of $s$ at $(a)\,\rho_P$=0.002 and $(b)\,\rho_P$=0.005.

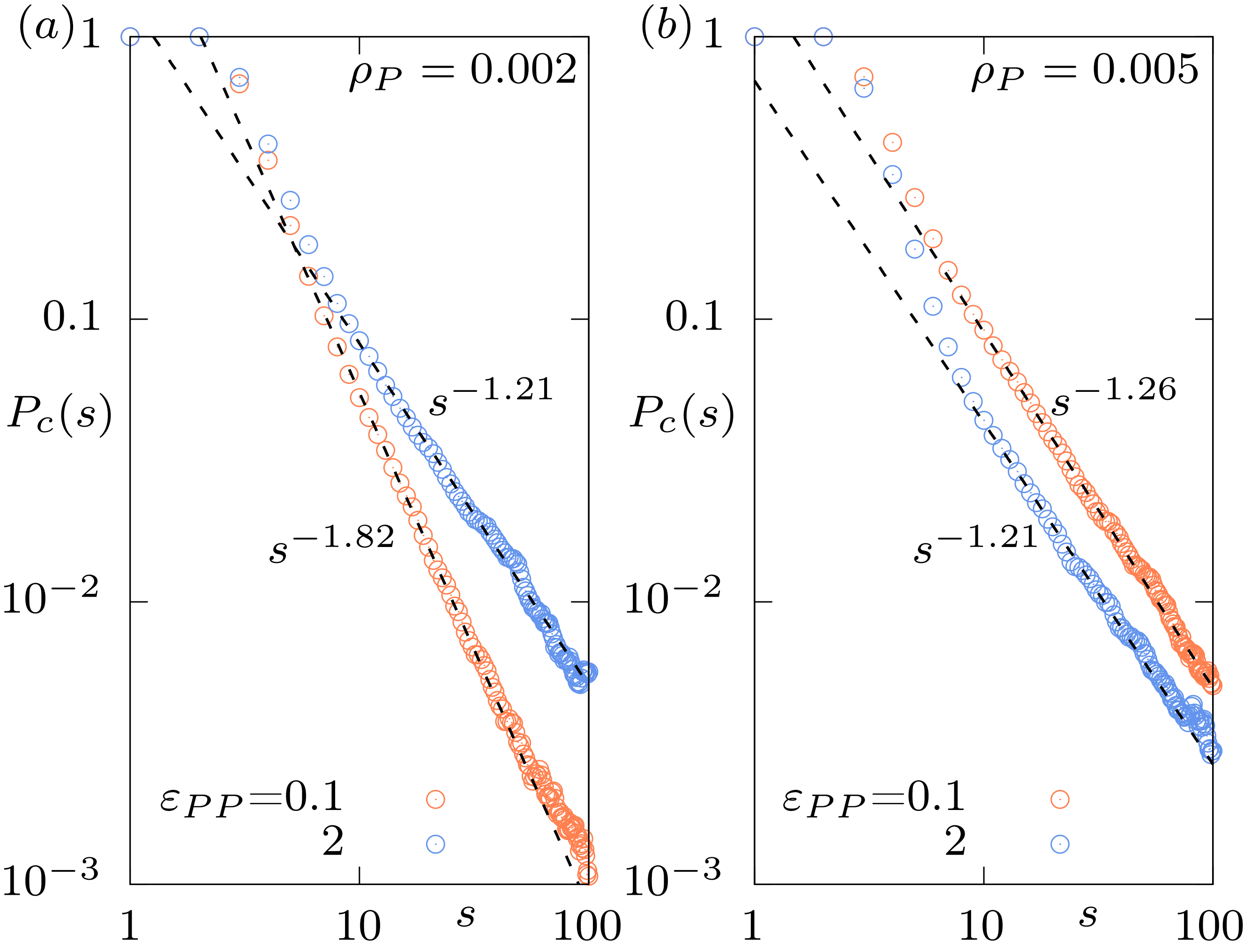


Fig. S5. Effect of $\varepsilon_{PP}$ on the contact probability between monomers of a homopolymer: $P_c(s)$ as a function of $s$ at $(a)$ $\rho_P$=0.002, and $(b)$ $\rho_P$=0.005.

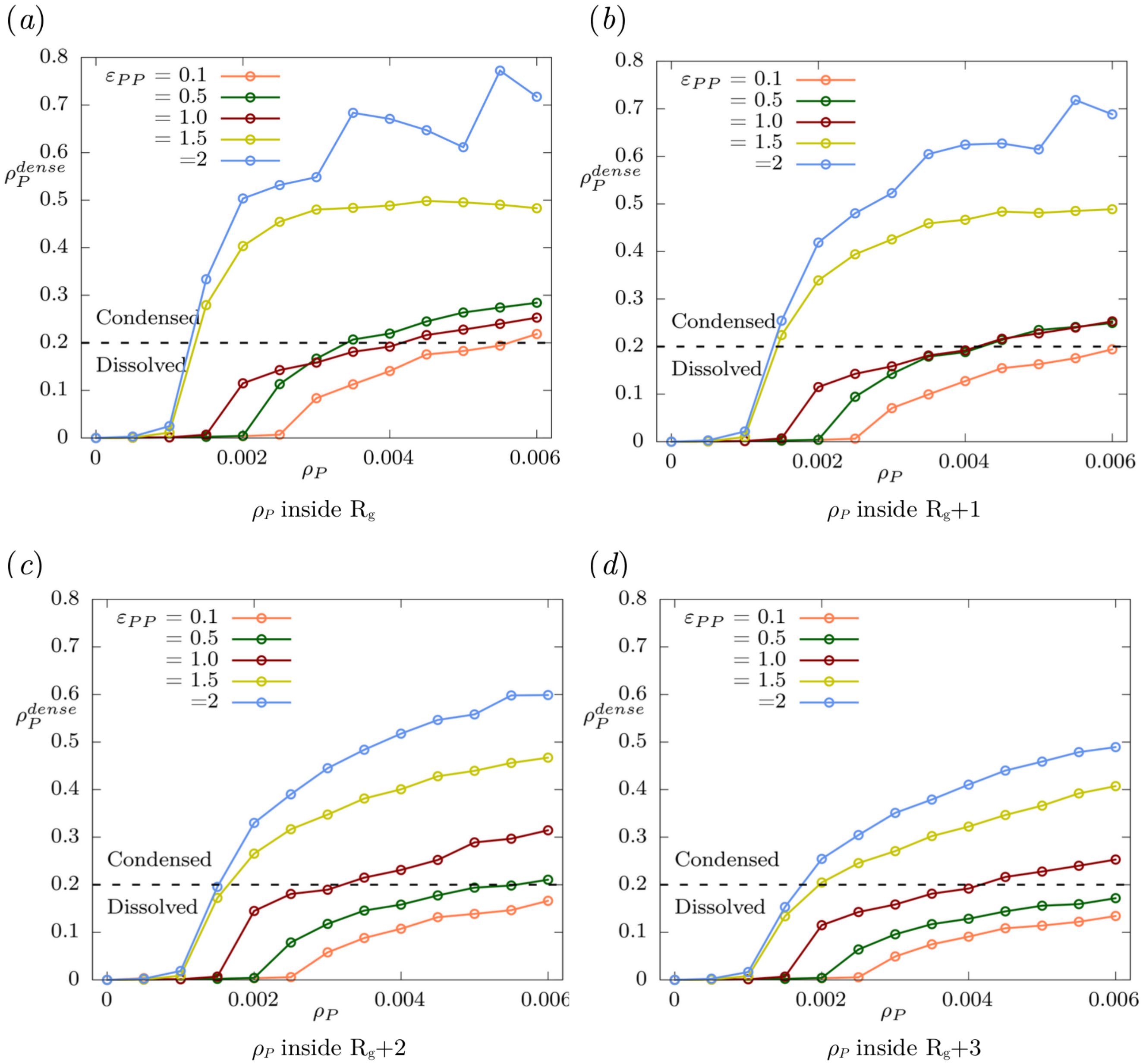


Fig. S6. Effect of increasing the thickness of the adsorbed layer ($\Delta r$ in the main text) on the density of condensed proteins ($\rho_P^{dense}$).

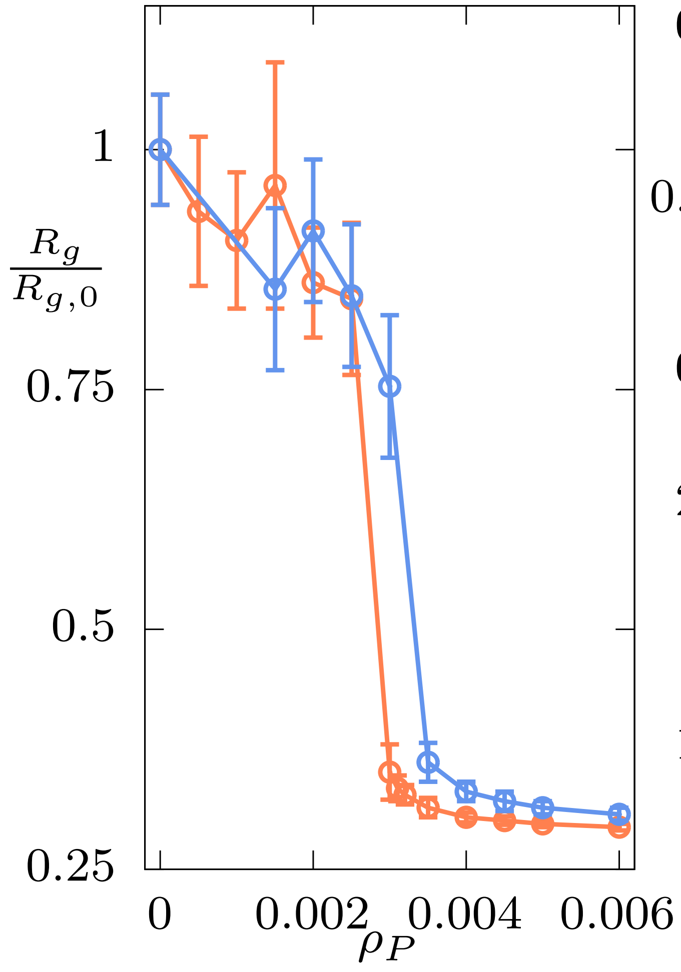


Fig. S7. $(a)$ Scaled radius of gyration $(R_g/R_{g,0})$, $(b)$ genomic loci exponent $(\nu)$, and $(c)$ contact probability exponent $(\alpha)$ of homopolymer with increase in $\rho_P$ at $\varepsilon_{PP}$=0.1 (attractive) in orange, and $\varepsilon_{PP}$=1 (repulsive) in blue.

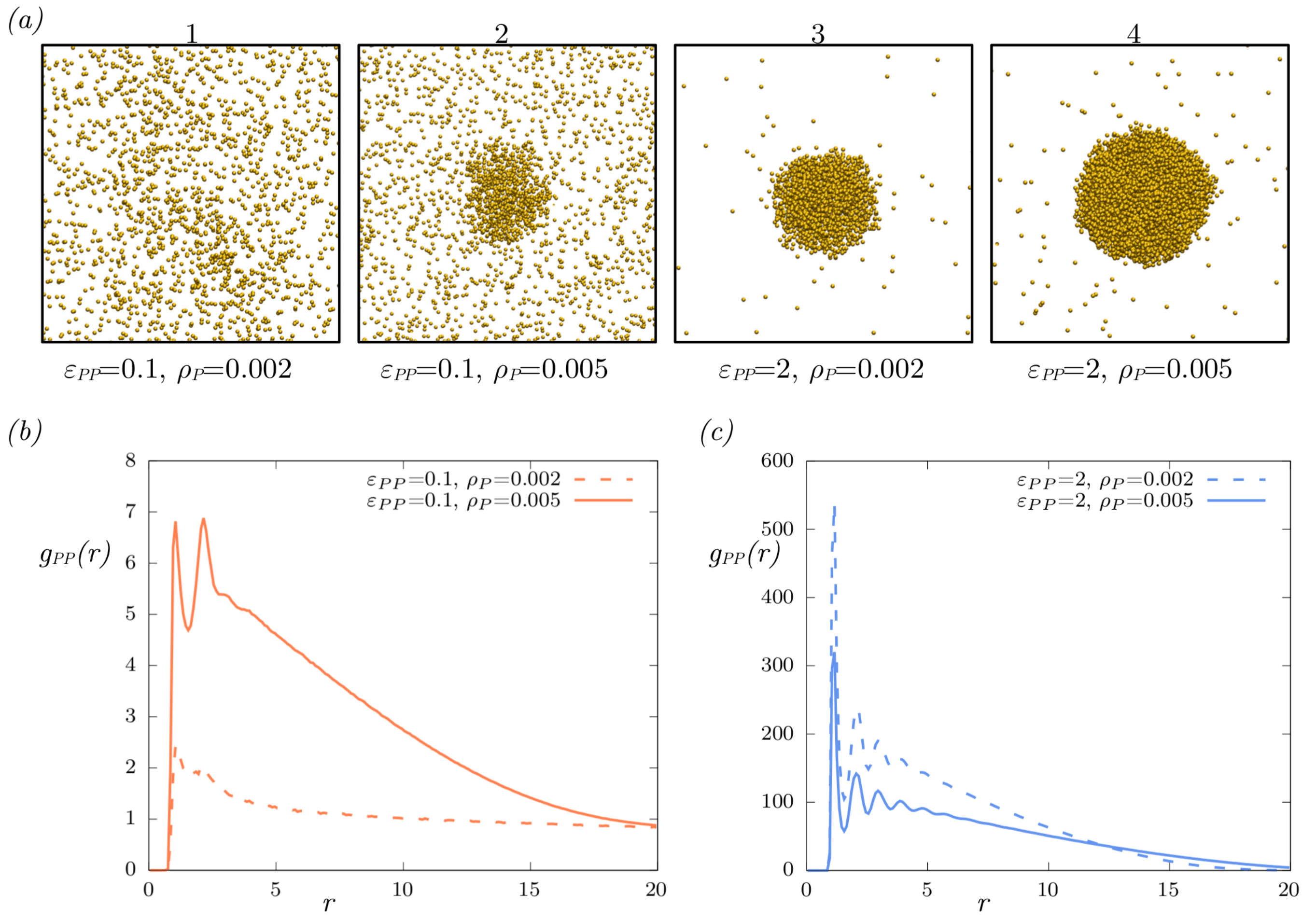


Fig. S8. $(a)$ Configuration of binders corresponding to Fig 2$(c)$ in the main text. $(b)$ and $(c)$ Radial distibution function between the binders ($g_{PP}(r)$) at $\varepsilon_{PP}$=0.1 and 2, respectively.

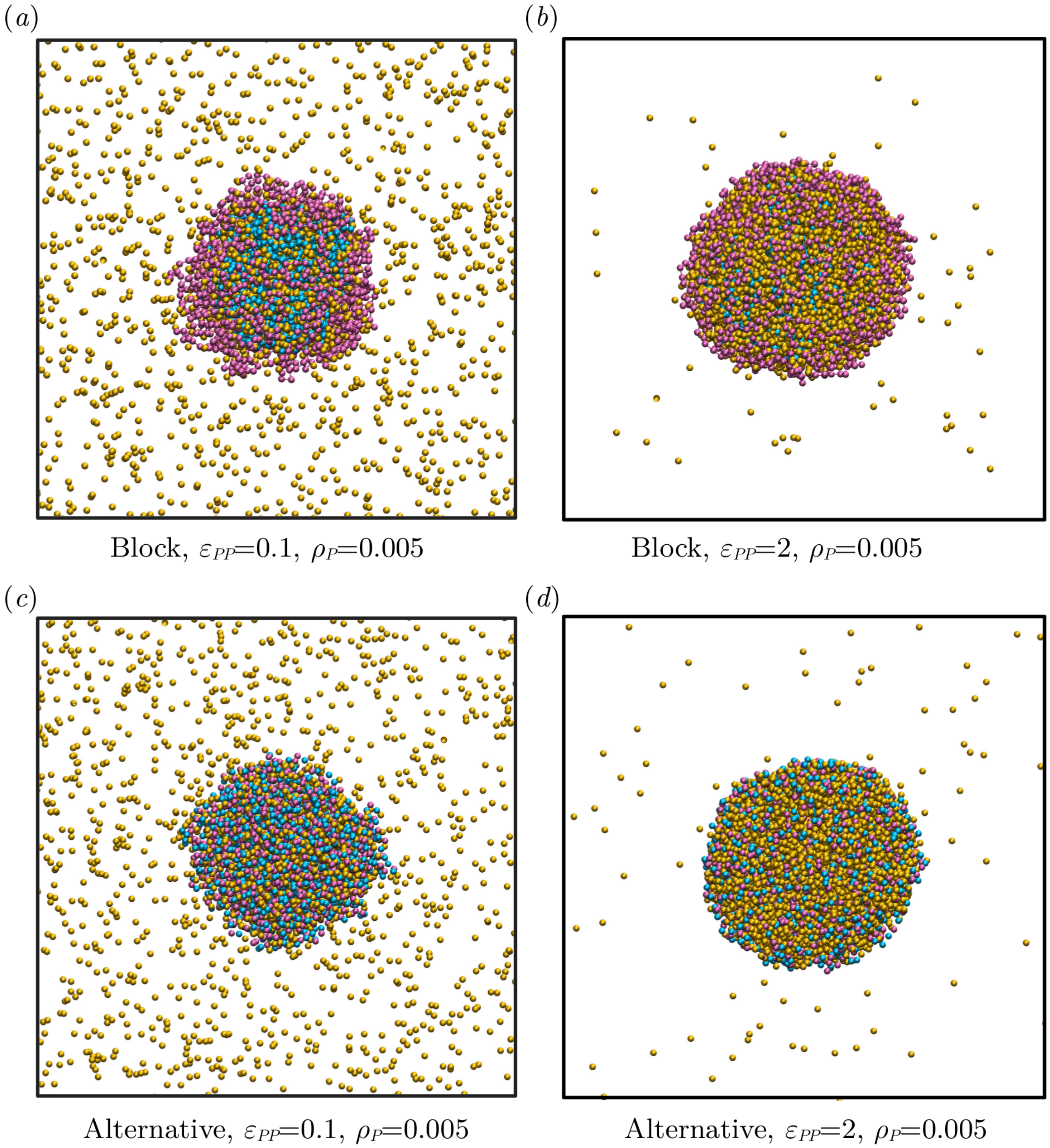


Fig. S9. Configurations of the block copolymer and alternative polymer along with the binders corresponding to Fig. 9 in the main text.

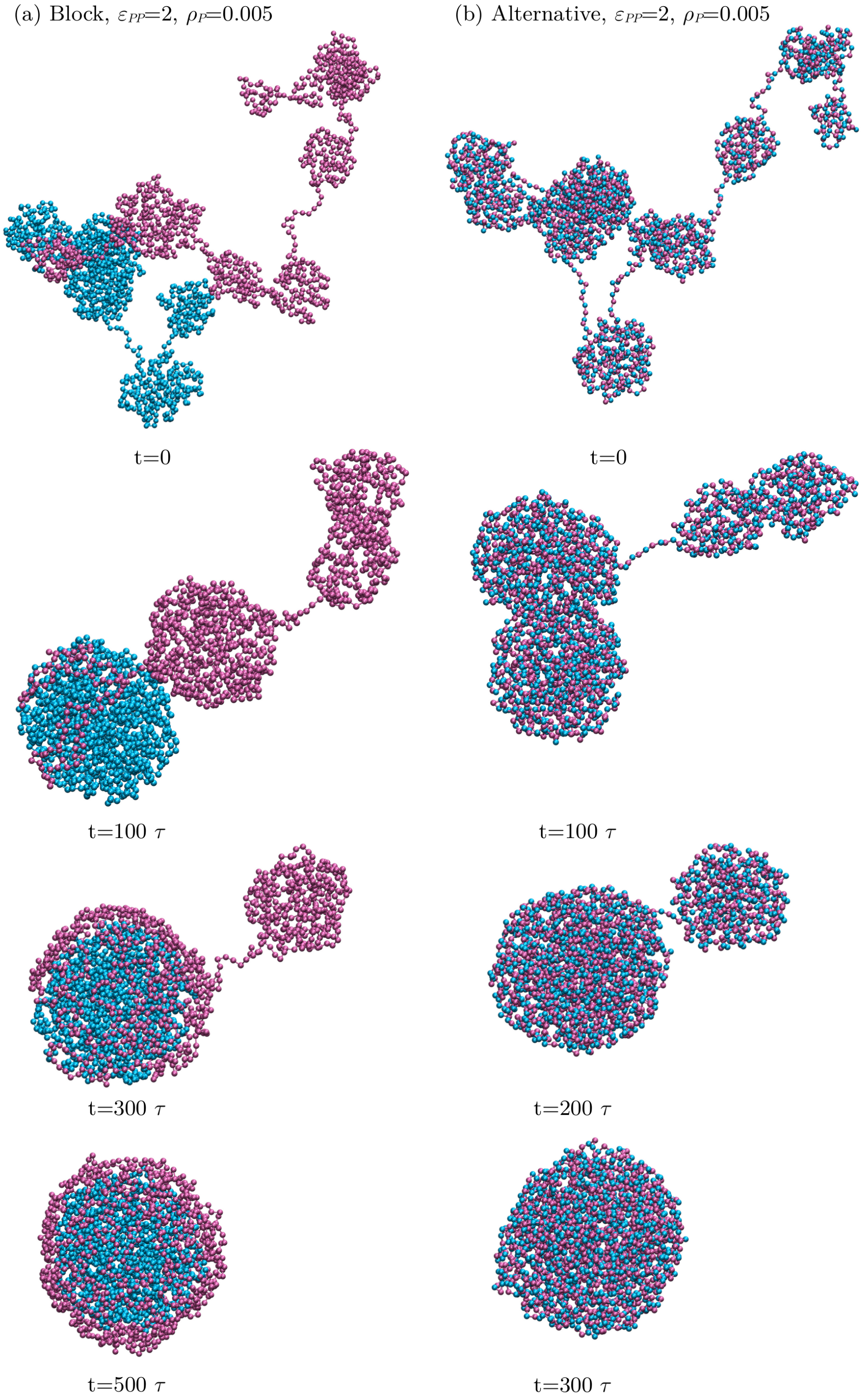


Fig. S10. Configurations of the block polymer (left column) and the alternative polymer (right column) showing the kinetics of collpase leads to layered organization in case of the former. We do not show the binders here for the purpose of clarity.
